# UTRGen: A unified framework for full-spectrum design of mRNA 5′ UTRs

**DOI:** 10.64898/2026.06.26.734691

**Authors:** Zhen Wang, Mingxuan Chen, Xiagu Zhu, Xingyu Fang, Zhaowen Cheng, Mei Lang, Junwei Zhang, Junjie Huang, Xiaolin Li

## Abstract

The 5′ untranslated region (5′ UTR) is a key regulatory element that governs mRNA translation and protein output. However, existing computational methods typically address isolated tasks such as functional prediction or sequence optimization, limiting their ability to support rational design across the full 5′ UTR engineering workflow. Here, we present UTRGen, a unified modeling framework for 5′ UTRs that integrates sequence generation, multi-property prediction, and constrained function-guided design. UTRGen is pre-trained autoregressively on large-scale 5′ UTR datasets from multiple species and subsequently adapted to diverse downstream regulatory tasks. Across systematic evaluations, UTRGen generates novel and diverse 5′ UTRs while preserving sequence, structural, and functional characteristics of natural UTRs. After task-specific fine-tuning, UTRGen achieves state-of-the-art performance across 14 benchmark datasets, improving translation efficiency prediction by up to 11.1%, expression level prediction by up to 13.2%, and mean ribosome load prediction by up to 3.0% relative to the strongest baselines. It also achieved the best overall performance for internal ribosome entry site identification. To enable controllable design, we formulate function-guided 5′ UTR design as a GRPO-based refinement process over a pre-trained autoregressive sequence prior, using composite rewards to encode functional objectives and biological constraints while regularizing toward the natural 5′ UTR distribution. The resulting sequences show consistently improved predicted translation efficiency and expression levels across cellular contexts, and reveal interpretable sequence features associated with high activity, including reduced C content, fewer upstream AUGs, and depletion of inhibitory motifs. Together, our results establish a unified modeling strategy for 5′ UTR design and lay a foundation for programmable control of translation.

## 1 Introduction

Messenger RNA (mRNA)-based therapeutics and vaccines have emerged as an important platform in modern medicine, but their efficacy depends critically on the stability and translational performance of exogenously delivered transcripts [1]. Among the cis-regulatory elements of mRNA that govern these properties, the 5′ untranslated region (5′ UTR) is particularly important because it directly regulates translation initiation and strongly influences protein output. Located upstream of the open reading frame, the 5′ UTR modulates ribosome recruitment and scanning, and can further affect mRNA stability, subcellular localization, and overall protein production [2]. In eukaryotic cells, the 5′ cap recruits the ribosome, which then scans along the 5′ UTR to recognize the start codon (AUG) and initiate translation [3]. Multiple features of the 5′ UTR, including its length, secondary structure, upstream open reading frames (uORFs), and specific sequence motifs, can affect ribosome loading and scanning efficiency, thereby shaping translation initiation efficiency [4]. Because 5′ UTR-mediated translational regulation is governed by multiple coupled sequence and structural determinants, its sequence-function relationships are difficult to resolve through conventional experimental characterization or heuristic rules alone. This complexity calls for more generalizable computational approaches that capture the multidimensional regulatory logic of 5′ UTRs.

This challenge has motivated the development of machine learning approaches for mapping 5′ UTR sequences to functional phenotypes. In particular, a variety of supervised learning models have been developed to predict mean ribosome load (MRL), translation efficiency (TE), expression level (EL), and internal ribosome entry site (IRES) activity from sequence information or manually engineered biological features [5, 6, 7, 8, 9]. These models have provided effective tools for evaluating 5′ UTR function and have advanced quantitative studies of translational regulation. However, such approaches typically rely on task-specific labeled data and often focus on a single functional phenotype, which limits their ability to support cross-task transfer, unified representation learning, and systematic modeling of multidimensional regulatory mechanisms. In recent years, inspired by the success of pre-trained language models in natural language processing, researchers have begun to introduce language modeling strategies into the study of mRNA, particularly 5′ UTR sequences, to learn generalizable representations from large-scale unlabeled data. Representative studies such as UTR-LM [10] and mRNABERT [11] have shown that Transformer-based pre-trained models can substantially improve predictive performance for tasks including MRL, TE, and EL, while also enhancing the representation of sequence semantics and regulatory features in mRNA. Although these studies have demonstrated the potential of pre-trained language models for functional analysis of mRNA, they have mainly focused on predicting the properties of existing sequences rather than enabling function-guided rational design, which offers a more direct route to generating 5′ UTRs for desired functional outcomes.

Designing 5′ UTRs to satisfy functional objectives is considerably more challenging than evaluating the properties of existing sequences, but it is also more closely aligned with the practical needs of mRNA engineering. Existing computational strategies for 5′ UTR design have often followed a generate-then-screen paradigm, in which deep generative models are used to produce candidate sequences and separate predictive models are subsequently applied for functional screening [12]. While this strategy has provided a useful starting point for computational 5′ UTR design, the separation between sequence generation and functional evaluation makes it less straightforward to guide design toward desired objectives directly. More broadly, existing studies have typically focused on individual components of the design process, such as sequence generation and function prediction, while lacking a unified framework capable of holistically supporting 5′ UTR generation, functional prediction, and function-guided design. However, in the context of therapeutic mRNA development, 5′ UTR design is inherently not a single-step problem but a systematic task encompassing sequence distribution learning, functional representation, multi-property prediction, and function-guided design. An ideal computational framework should not only be able to learn the sequence patterns of natural 5′ UTRs and generate biologically plausible candidate sequences, but also provide unified modeling and accurate prediction of multiple functional properties, while further supporting controllable design toward specific performance objectives.

Based on these considerations, we developed UTRGen, a unified framework for full-spectrum modeling and design of mRNA 5′ UTRs. UTRGen is designed to support three tightly connected capabilities: (i) generation of 5′ UTR sequences that follow the distributional patterns of natural sequences, (ii) prediction of multiple functional properties of 5′ UTRs, and (iii) constrained design of 5′ UTRs toward specified functional objectives. Specifically, UTRGen is pre-trained autoregressively on a diverse, high-quality dataset containing approximately 9.45 million non-redundant 5′ UTR sequences from 573 eukaryotic species, systematically curated and constructed by the authors, with details provided in the Methods section (Fig. 1a). Through large-scale cross-species pre-training, the model captures conserved and transferable regulatory patterns in 5′ UTR sequences, enabling the generation of novel sequences that remain consistent with natural distributional properties while also providing information-rich representations for downstream tasks (Fig. 1a). Built on this pre-training foundation, UTRGen further supports unified prediction of multiple translation-related phenotypes, including MRL, TE, EL, and IRES activity (Fig. 1b). To overcome the limitations of existing methods in achieving function-guided design, we introduced a reinforcement learning strategy based on group relative policy optimization (GRPO) [13, 14] on top of the pre-trained generative model to enable function-guided design of 5′ UTRs (Fig. 1c). In this process, model optimization is driven by a composite reward mechanism. The reward incorporates functional objective signals from the predictor, particularly TE or EL. These two properties were selected as the primary optimization targets because they provide complementary and application-relevant readouts of 5′ UTR function, reflecting translational efficiency and overall expression output, respectively. The reward also includes explicit penalty terms and constraints based on factors such as GC content and minimum free energy (MFE). In addition, we introduced a Kullback-Leibler (KL) regularization term to constrain the optimized sequences to remain as close as possible to the natural sequence distribution learned during pre-training. This design not only improves the biological plausibility and diversity of the generated sequences but also reduces the risk of degenerate solutions. Taken together, these modules enable UTRGen to efficiently explore the broad 5′ UTR sequence space and generate candidate sequences that satisfy both design objectives and biological constraints.

**Figure 1.**
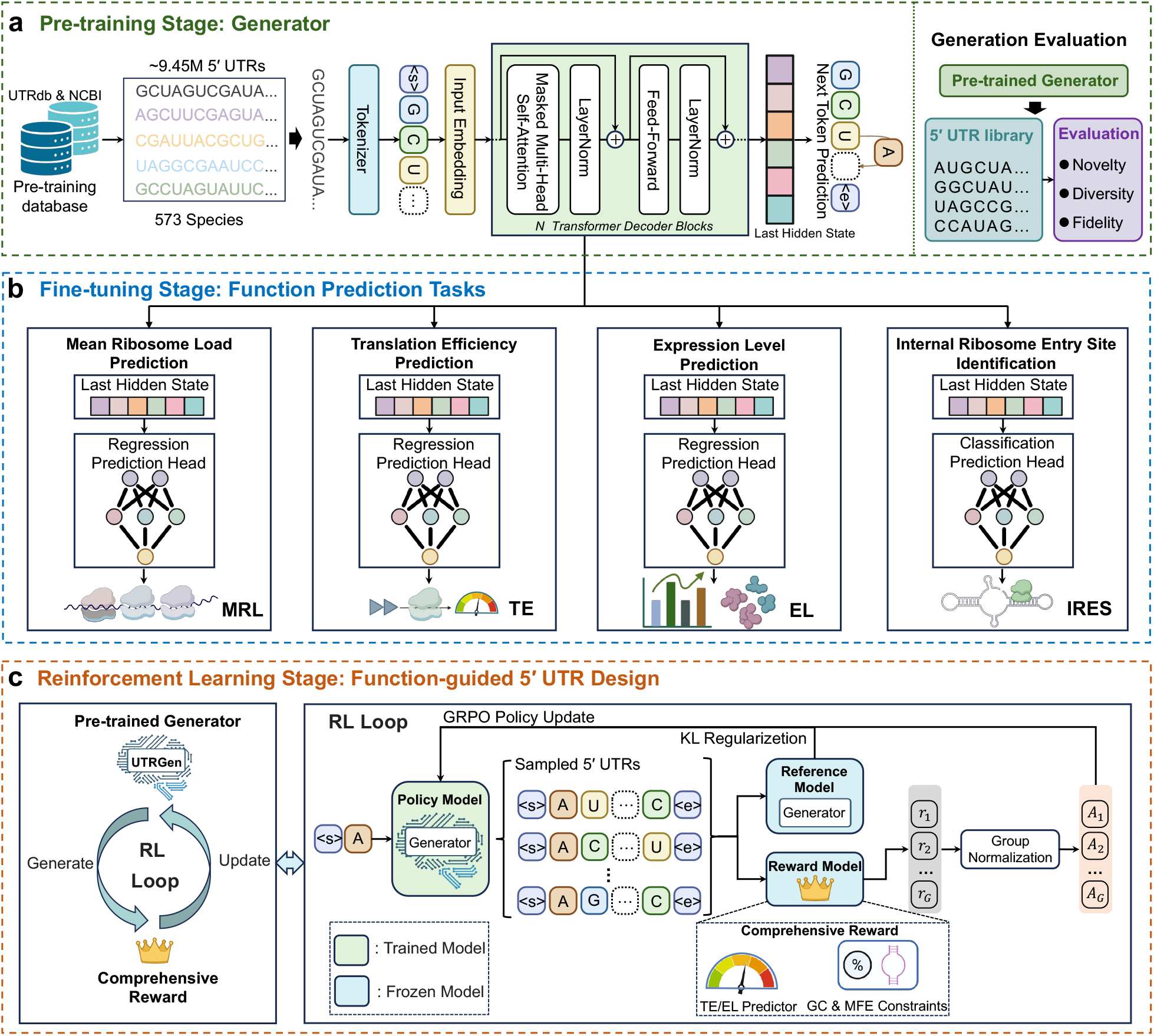
Overview of the UTRGen framework. **a**, Autoregressive pre-training of UTRGen as a 5′ UTR generator on ∼ 9.45 million sequences from 573 eukaryotic species (UTRdb and NCBI) using next-token prediction, followed by generation-level evaluation of sampled 5′ UTR libraries (novelty, diversity, and fidelity). **b**, Fine-tuning of the pre-trained backbone for downstream function prediction by attaching task-specific heads to the last hidden state to predict MRL, TE, EL, and IRES activity. **c**, Reinforcement learning for function-guided 5′ UTR design. A GRPO policy is optimized from the pre-trained generator using a composite reward (TE/EL predictor with additional GC and MFE constraints) and KL regularization to a reference model, enabling controllable exploration of the 5′ UTR design space. Icons in figures were created with BioRender.com.

We systematically evaluated UTRGen across sequence generation, functional prediction, and function-guided design, demonstrating integrated capabilities that span the full workflow of 5′ UTR design. In sequence generation, UTRGen produced novel and diverse 5′ UTR sequences that remained highly consistent with natural sequences across multiple distributional, structural, and functional metrics. In benchmark tests for 5′ UTR functional prediction, UTRGen showed state-of-the-art performance across 14 datasets spanning four task categories: MRL, TE, EL, and IRES prediction. Relative to the strongest baselines, UTRGen improved performance by up to 11.1% for TE prediction, 13.2% for EL prediction, and 3.0% for MRL prediction, and also delivered the best overall performance for IRES identification. In function-guided design, UTRGen generated candidate 5′ UTRs with consistently improved predicted TE and EL across multiple cell-line contexts.

In addition, we performed a systematic analysis of key sequence features and regulatory motifs in the RL-designed sequences. The results showed that, across different cellular contexts, the model could spontaneously learn interpretable sequence characteristics associated with high TE, including reduced C content and systematic avoidance of known inhibitory elements. Specifically, as the predicted TE or EL score increased, the average C content of the generated sequences showed a decreasing trend, a finding that aligns with previous research [15]. At the same time, upstream start codons (uAUGs) [16] and several previously reported translation-repressive motifs [17] were significantly depleted in the RL-designed sequences. These results indicate that UTRGen not only enables function-guided design but also captures composition constraints and sequence patterns consistent with translational regulation without explicit encoding of biological rules, thereby providing interpretability for its design outcomes. In summary, UTRGen provides a unified computational framework that organically integrates 5′ UTR sequence generation, multi-property prediction, and function-guided design, offering a more systematic and flexible methodological foundation for full-spectrum design of 5′ UTRs and extending their potential applications in therapeutic mRNA engineering.

## 2 Results

### 2.1 Overview of the UTRGen and benchmarks

UTRGen adopts a GPT-style autoregressive pre-training architecture built on a Transformer decoder. This generative formulation enables two complementary capabilities within a unified framework: (i) learning the statistical regularities of 5′ UTR sequences to derive informative functional representations for diverse downstream prediction tasks; and (ii) proposing candidate sequences in the vast design space via token-by-token generation, providing an actionable generator for subsequent constraint-aware optimization and function-guided design. As summarized in Fig. 1, our framework comprises three stages.

#### Stage I: autoregressive pre-training for 5′UTR generation

We pre-trained UTRGen using approximately 9.45 million 5′ UTR sequences, covering 573 eukaryotic species, compiled from UTRdb [18] and NCBI [19]. Using a next-token prediction (NTP) objective, the model learned nucleotide-level sequence regularities and long-range dependencies, with the pre-training loss decreasing steadily and converging over the course of training (Supplementary Fig. 1), thereby yielding a general-purpose generator. We further assessed the generative behaviour of the pre-trained model by sampling 5′ UTR libraries and quantifying novelty, diversity, and fidelity (Fig. 1a).

#### Stage II: fine-tuning for function prediction

Starting from the pre-trained backbone, we attach lightweight task-specific heads to the last hidden state and fine-tune UTRGen to predict multiple translation-related phenotypes (Fig. 1b). These tasks include MRL, TE, EL, and IRES activity. This stage aligns the unsupervised sequence prior with supervised functional signals, yielding quantitatively comparable predictors that can serve as scoring functions for downstream design.

#### Stage III: reinforcement learning for function-guided 5′UTR design

Sampling from the pre-trained generator alone does not reliably produce sequences that satisfy engineering objectives. We therefore perform reinforcement learning with GRPO, initializing the policy from the pre-trained UTRGen generator and using the fine-tuned TE/EL predictor as the primary reward signal (Fig. 1c). To support controllable exploration, we define a composite reward that augments TE/EL with explicit constraint terms, including GC content and MFE. During policy updates, we apply KL regularization to a reference generator to keep the optimized policy close to the natural 5′ UTR distribution captured during pre-training, thereby improving sequence plausibility and diversity and reducing reward hacking. This closed-loop optimization progressively shifts the generator towards desired translation phenotypes and yields novel candidate 5′ UTRs while maintaining sequence plausibility.

To evaluate downstream predictive performance, we benchmarked UTRGen against three categories of representative methods, including conventional neural network baselines, general-purpose ncRNA pre-trained language models, and mRNA-specific pre-trained models, across diverse tasks and data regimes. Specifically, for MRL prediction across 10 datasets, we compared UTRGen with 13 baselines, including neural network approaches Optimus [5], FramePool [8], and MTtrans [7]; general ncRNA language models RNABERT [20], RNA-FM [21], RNAErnie [22], ERNIE-RNA [23], and RiNALMo [24]; and mRNA-specific language models CaLM [25], mRNAFM (an mRNA-pretrained extension of RNA-FM) [21], UTR-LM [10], mRNABERT [11], and GEMORNA-UTR [12]. For TE and EL prediction across three datasets, we benchmarked against 10 methods, covering neural network baselines Optimus [5], Cao-RF [6], FramePool [8], and MTtrans [7]; general ncRNA language models RNABERT [20] and RNA-FM [21]; and mRNA-specific models mRNA2vec [26], UTR-LM [10], mRNABERT [11], and GEMORNA-UTR [12], thereby testing effectiveness on translation- and expression-relevant phenotypes closer to practical applications. Finally, for IRES activity prediction, we compared against six representative methods—IRESpy [27], IRESfinder [28], FramePool [8], UTR-LM [10], mRNABERT [11], and GEMORNA-UTR [12]—to examine the model’s ability to capture signals related to non-canonical translation initiation and to further assess transferability when combining pre-trained representations with task supervision.

### 2.2 UTRGen generates novel and diverse 5′UTRs while preserving native-like sequence properties

To evaluate the foundational sequence modeling capacity conferred by large-scale pre-training, we first examined the base generative model trained solely through self-supervised learning. An ideal biological sequence generator should not only accurately capture the underlying distribution of native sequences, but also be capable of exploring previously uncharacterized regions of sequence space. On this basis, we systematically evaluated the generated 5′ UTRs from three perspectives: novelty, diversity, and fidelity. Across four length intervals (25-50 nt, 50-100 nt, 100-150 nt, and 150-200 nt), we generated 10,000 unique sequences for each bin, yielding a total of 40,000 generated sequences (Generated) for analysis. In parallel, we sampled 10,000 unique sequences from both the pre-training training set (Training) and test set (Test) for each length interval, resulting in 40,000 native sequences from each set as controls (details are provided in Supplementary Note 1.2).

To assess the novelty of UTRGen-generated sequences and determine whether they were simply reproductions of training examples, we performed sequence alignments by comparing the generated and test sets against the training set using BLAT [29]. The generated sequences showed minimal sequence identity to the training set, with only 4.53% having at least one alignment hit of ≥ 20 nt (Fig. 2a), and the longest match lengths were all below 50 nt, with a median of only 27 nt (Fig. 2b). These results indicate that UTRGen does not merely memorize training examples but can generate previously unseen 5′ UTR sequences. In addition, the generated sequences exhibited high internal diversity. The internal alignment match rate within the generated set (17.71%) was higher than that within either the training or test set (Fig. 2c), consistent with the expectation that the model had learned recurrent local sequence patterns present in native 5′ UTRs. Nevertheless, the median of the longest self-alignment match length between generated sequences was only 41 nt (Fig. 2d). Furthermore, among the generated sequences, which had an average length of 124.2 nt, the average pairwise Levenshtein distance reached 98.25 (Supplementary Fig. 2a), indicating substantial sequence-level differences at the global scale. Collectively, these results show that UTRGen can generate highly diverse sequences without copying the training data.

**Figure 2.**
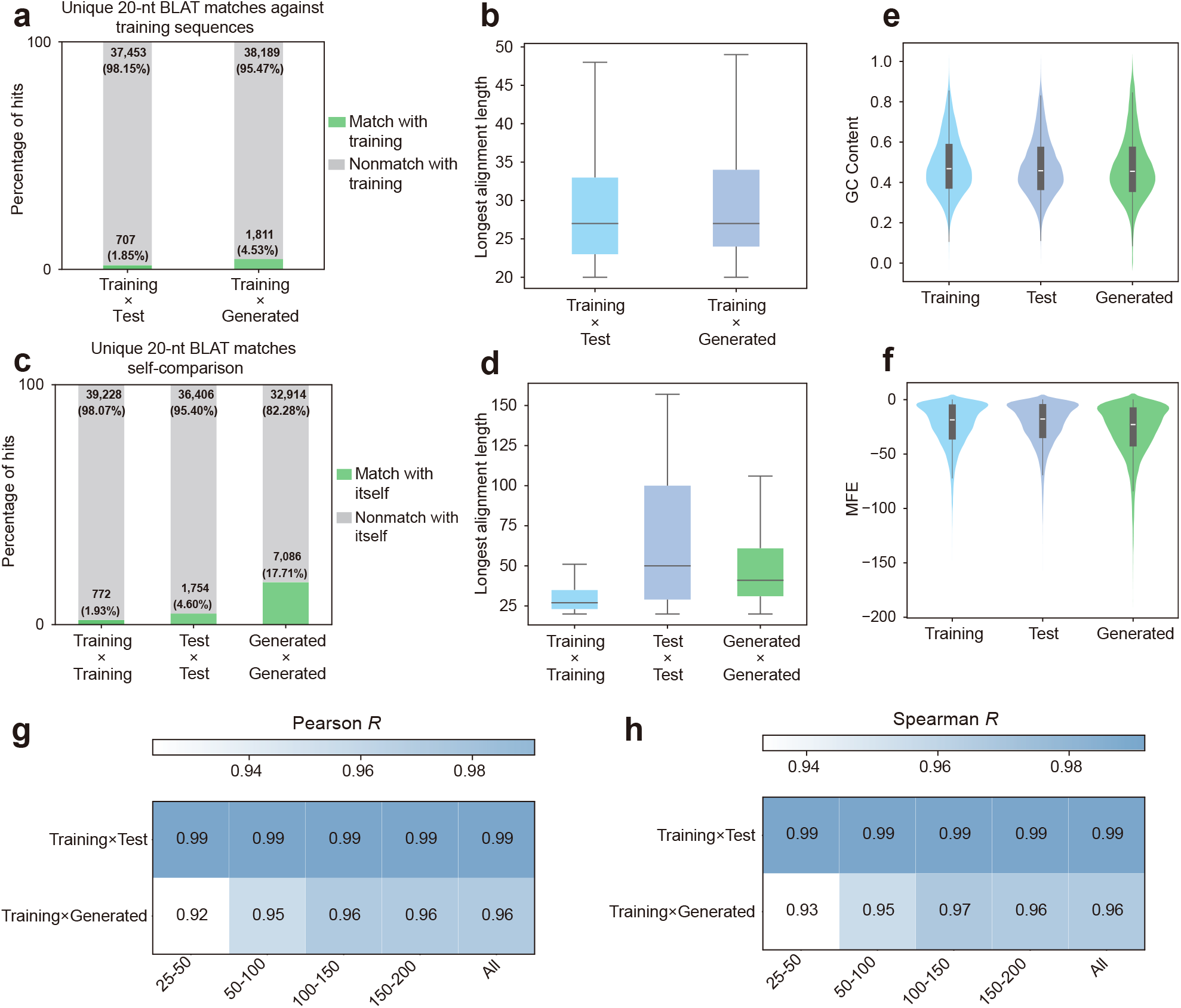
UTRGen generates novel, diverse, and native-like 5′ UTR sequences. **a**, The hit rate for UTRGen-generated sequences and Test set sequences finding at least a 20-nucleotide match in the Training set. The majority (95.47%) of UTRGen-generated sequences do not find any match in the Training set. **b**, Distribution of the longest match length between the Generated set and the Test set sequences against the Training set. **c**, The self-alignment hit rates for the Generated set, Test set, and Training set. **d**, Distribution of the longest self-alignment match length for the Generated set, Test set, and Training set. **e, f**, The GC content **(e)** and MFE **(f)** distributions for Training set, Test set, and Generated set, demonstrating high compositional and thermodynamic stability. **g, h**, Heatmaps showing Pearson **(g)** and Spearman **(h)** correlation coefficients of 4-mer frequencies. For four different length intervals and overall data (All), the Training set is compared separately with the Test set and the Generated set.

Having confirmed novelty and diversity, we assessed whether the generated sequences captured key properties of native 5′ UTRs. The distributions of GC content (Fig. 2e) and MFE (Fig. 2f) in the generated sequences closely resembled those of native 5′ UTRs, suggesting that UTRGen captures constraints related to both sequence composition and secondary structure. To further assess local sequence grammar, we compared 4-mer frequency distributions between generated and native sequences. Because 4-mer features reflect short-range sequence composition patterns and are closely associated with non-coding RNA analysis and gene expression regulation [30], they provide a useful measure of local sequence organization. As shown in Fig. 2g and h, the 4-mer frequency distributions of generated sequences were highly concordant with those of the training set across all length intervals, with both Pearson and Spearman correlation coefficients exceeding 0.9 in every case, indicating that UTRGen accurately learned representative local patterns of 5′ UTR sequences. We further examined the relationship between generated and native sequences in the latent space. UTRGen-generated sequences broadly and uniformly covered the regions occupied by native sequences, while showing a modest outward extension at the boundaries (Supplementary Fig. 2b). This result suggests that UTRGen preserves the global distributional structure of native 5′ UTRs while retaining the ability to explore plausible novel sequence variants. Taken together, large-scale self-supervised pre-training provides UTRGen with a strong generative prior, enabling it to produce 5′ UTR sequences that are novel, diverse, and native-like in their sequence properties.

### 2.3 UTRGen achieves state-of-the-art prediction of mean ribosome load

Ribosome load, defined as the number of ribosomes bound to an individual mRNA molecule at any given time, is a key determinant of protein synthesis efficiency. It provides a quantitative readout of translation initiation and serves as a proxy for expected protein expression. Accordingly, accurately predicting ribosome load from 5′ UTR sequences alone is critical for optimizing mRNA sequence design toward maximal protein output, particularly when constructing novel 5′ UTRs beyond the distribution of existing templates.

To address this challenge, we leveraged a benchmark dataset established by prior studies using massively parallel reporter assays (MPRA) [5]. The dataset comprises eight synthetic 5′ UTR libraries of fixed length (50 nt), spanning distinct uridine-modification and reporter conditions (U_1_, U_2_, Ψ_1_, Ψ_2_, m1Ψ_1_, m1Ψ_2_, mC-U_1_ and mC-U_2_); each library contains approximately 280,000 sequences paired with their corresponding ribosome-load measurements. In this work, we initialized our model with the pre-trained UTRGen and subsequently performed supervised fine-tuning to predict biological functions, such as MRL. Specifically, we extracted nucleotide-level hidden states from the final layer of the pre-trained decoder and fed them into a downstream prediction head (1D-CNN) (Fig. 3a).

**Figure 3.**
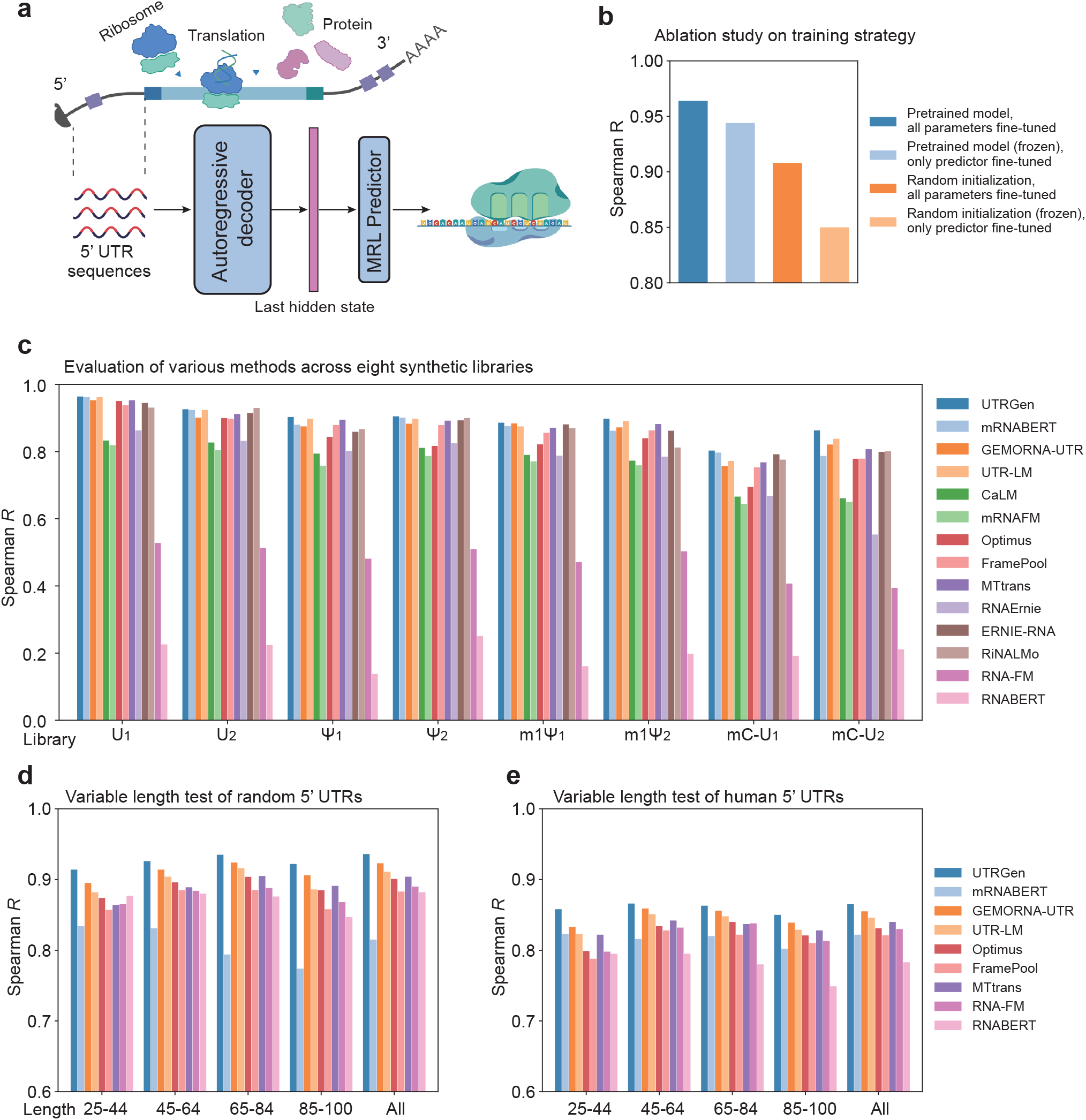
Comparison of MRL prediction tasks. **a**, Schematic of the UTRGen architecture for downstream function prediction. Upon receiving a 5′ UTR sequence, the model extracts hidden states from the final autoregressive decoder layer and routes them to a dedicated predictor to generate functional metrics, such as MRL. **b**, Ablation study of training strategies. MRL prediction performance is compared between pre-trained and randomly initialized weights, under both all-parameter fine-tuning and frozen-backbone (prediction head only) configurations. **c**, Benchmarking of UTRGen against 13 baseline models across eight independent fixed-length (50 nt) synthetic 5′ UTR libraries. **d, e**, Evaluation of accuracy and generalization on variable-length 5′ UTR datasets. Performance is assessed on a randomly synthesized test set (25-100 nt) **(d)** and a real human endogenous 5′ UTR dataset **(e)**.All results are stratified by sequence length intervals. Icons in figures were created with BioRender.com.

To quantify the benefit of pre-training, we compared the “pre-training and all-parameter fine-tuning” strategy against alternative settings, including training without pre-trained weights and schemes with parameter freezing or partial fine-tuning. As shown in Fig. 3b, pre-training followed by all-parameter fine-tuning achieved the best predictive performance and significantly outperformed the other configurations, supporting the effectiveness of our pre-training and fine-tuning strategy.

We next performed a systematic comparison of UTRGen against 13 baseline models, including task-tailored machine-learning approaches (e.g., Optimus [5], FramePool [8], and MTtrans [7]), general ncRNA language models (e.g., RNA-FM [21], RNAErnie [22], ERNIE-RNA [23], and RiNALMo [24]), and mRNA-specific language models (e.g., UTR-LM [10], GEMORNA-UTR [12], and mRNABERT [11]). Given that GEMORNA-UTR [12] was not originally developed for MRL prediction, we retrained it on the MRL datasets under the same evaluation setting; the remaining baseline results were taken from the publicly available mRNABERT [11] report and follow the evaluation protocol described therein. Across the eight synthetic libraries, UTRGen consistently delivered top-tier prediction accuracy, ranking first in seven of the eight libraries and second in the remaining library (Fig. 3c and Supplementary Table 1). Relative to the strongest baseline in each library, UTRGen improved the Spearman *R* score by up to 3.0%, with the largest gains observed on modified UTR libraries.

To further assess generalization to 5′ UTRs of varying lengths, we evaluated UTRGen on two independent variable-length datasets introduced by Sample *et al*. [5] and widely adopted in subsequent studies as benchmarks. These datasets comprise synthetic and human 5′ UTRs with lengths ranging from 25 to 100 nt. Further details are provided in Methods. Following the length-stratified held-out testing protocol used by Optimus [5] and UTR-LM [10], we fine-tuned the pre-trained model on 76,319 random 5′ UTR sequences (25–100 nt), and then evaluated it on 7,600 random and 7,600 human 5′ UTRs alongside the baseline methods (Fig. 3d, e, and Supplementary Table 2). UTRGen achieved the best overall performance on both random and human 5′ UTR benchmarks, with Spearman *R* scores of 0.935 and 0.862, respectively. Relative to the strongest baseline, this corresponds to improvements of 1.3% on the random dataset and 0.8% on the human dataset. UTRGen also ranked first in all random length strata and in three of the four human length strata, indicating strong robustness to length variation and effective transfer across distinct 5′ UTR contexts.

### 2.4 UTRGen improves prediction of translation efficiency and expression level

Protein output is governed primarily by two processes, transcription and translation. Under steady-state conditions, protein abundance largely depends on mRNA EL and transcript TE. EL quantifies the relative abundance of cellular transcripts and is measured using RNA-seq RPKM, where RPKM denotes the number of reads per kilobase of transcript per million mapped reads [6]. TE captures how efficiently an mRNA is translated and is conventionally defined as the ratio of Ribo-seq RPKM (reflecting ribosome-protected footprints) to RNA-seq RPKM [6].

In this section, we used three endogenous benchmark datasets (constructed by Cao *et al*. [6] and widely adopted in subsequent studies), derived from human muscle tissue (muscle), the human prostate cancer cell line PC3 (PC3), and the human embryonic kidney (HEK) 293T cell line. Following the setup of Chu *et al*. [10], we performed supervised fine-tuning of the pre-trained UTRGen model with ten-fold cross-validation to predict TE and EL. To quantify the contribution of pre-training, we further conducted ablation experiments on the muscle dataset. The results show that, compared with random initialization or freezing/partial fine-tuning alternatives, pre-training initialization followed by full-parameter fine-tuning yields the best performance on both TE and EL prediction tasks (Fig. 4a, b), supporting the effectiveness of this training paradigm on endogenous data.

**Figure 4.**
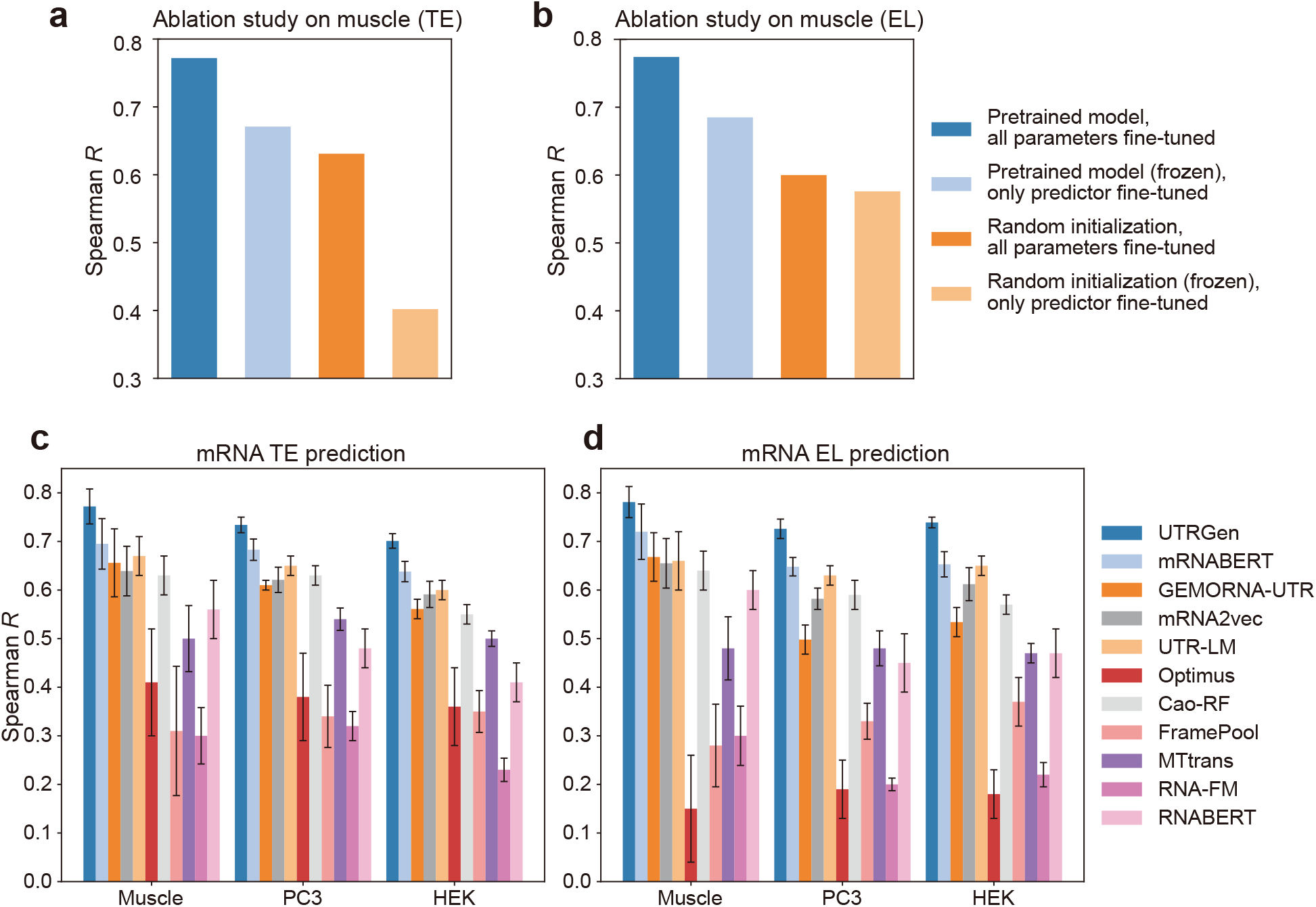
Prediction of mRNA TE and EL for endogenous datasets. **a, b**, Ablation of training strategies on the muscle dataset. Spearman *R* for TE **(a)** and EL **(b)** is shown for pre-trained versus randomly initialized models, under all-parameter fine-tuning and frozen-backbone (head-only) training. **c, d**, Comparison with 10 baselines for TE (**c**) and EL (**d**) prediction across Muscle, PC3 and HEK datasets. Bar heights represent mean Spearman *R* values from ten-fold cross-validation, with error bars indicating s.d. Statistical significance was assessed by paired *t*-tests on fold-wise results (Supplementary Table 4). UTRGen achieved the highest mean Spearman *R* in all six dataset-task combinations and significantly outperformed the best-performing baseline in each setting (*P <* 0.05).

For a systematic benchmark, we compared UTRGen with ten established baselines, including mRNABERT [11], GEMORNA-UTR [12], mRNA2vec [26], UTR-LM [10], Optimus [5], Cao-RF [6], FramePool [8], MTtrans [7], RNA-FM [21], and RNABERT [20]. Because mRNABERT and GEMORNA-UTR were not developed for TE/EL prediction, and because the original mRNA2vec study did not report results under ten-fold cross-validation, we retrained these models on the same TE/EL datasets under the same ten-fold cross-validation protocol to ensure a fair comparison; results for the remaining baselines were taken from the publicly available UTR-LM report and follow the evaluation protocol described therein.

As shown in Fig. 4c, d, and Supplementary Table 3, UTRGen achieved the best performance for both TE and EL prediction across all three datasets. Paired *t*-tests on fold-wise results showed that these improvements over the best-performing baseline were statistically significant in all six TE/EL benchmark settings (Supplementary Table 4). In particular, compared with mRNABERT, which was the strongest baseline across all six settings, UTRGen improved TE prediction by 11.1%, 7.5%, and 9.9% on Muscle, PC3, and HEK, respectively, and EL prediction by 8.5%, 12.0%, and 13.2% on the same datasets. Relative to UTR-LM, UTRGen improved TE prediction by 15.2%, 12.9%, and 16.8% across the three datasets, and EL prediction by 18.3%, 15.2%, and 13.7%. Relative to GEMORNA-UTR, the corresponding improvements were 17.7%, 20.3%, and 25.0% for TE, and 16.9%, 45.8%, and 38.4% for EL. Together, these results indicate that UTRGen effectively captures sequence features associated with endogenous transcript abundance and translational efficiency across diverse tissue and cellular contexts.

### 2.5 UTRGen enables accurate identification of internal ribosome entry sites

IRESs are cis-acting RNA elements that are typically located in the 5′ UTR of mRNAs. In contrast to the canonical cap-dependent initiation mechanism, which relies on the 5′ cap and scanning from the transcript 5′ end, IRESs can directly recruit ribosomes and associated initiation factors at internal positions to drive cap-independent translation. This mechanism is particularly important for viral mRNAs that lack a 5′ cap, and it also contributes to the selective translation of specific cellular transcripts under stress and other conditions. Although it has been estimated that 10% of cellular and viral mRNAs may initiate translation via IRESs, the number of experimentally validated and systematically annotated IRESs remains limited. As a result, predictors that rely on heuristic rules or handcrafted feature engineering (for example, IRESPred [31], IRESfinder [28], and IRESpy [27]) often fail to produce consistent and reliable predictions across datasets and experimental contexts. Therefore, accurate identification of putative IRES elements within 5′ UTRs is essential not only for elucidating the molecular basis of cap-independent translation, but also for enabling mRNA design and optimization for expression control.

To evaluate whether UTRGen captures sequence determinants of IRES activity, we fine-tuned the pre-trained model as a binary classifier by attaching a task-specific classification head to the backbone representations (Fig. 5a). We constructed an IRES benchmark dataset by integrating sequences from multiple resources, including IRESbase [32], Rfam [33], IRESite [34], IRESPred [31], Weingarten-Gabbay *et al*. [35], and Chen *et al*. [36], followed by unified filtering and deduplication across sources. The resulting dataset contained 51,053 sequences, including 25,210 IRES-positive sequences and 25,843 non-IRES sequences, with average lengths of approximately 190 nt and 179 nt, respectively (Fig. 5b). This provided a relatively balanced benchmark for IRES prediction (see Methods for details).

**Figure 5.**
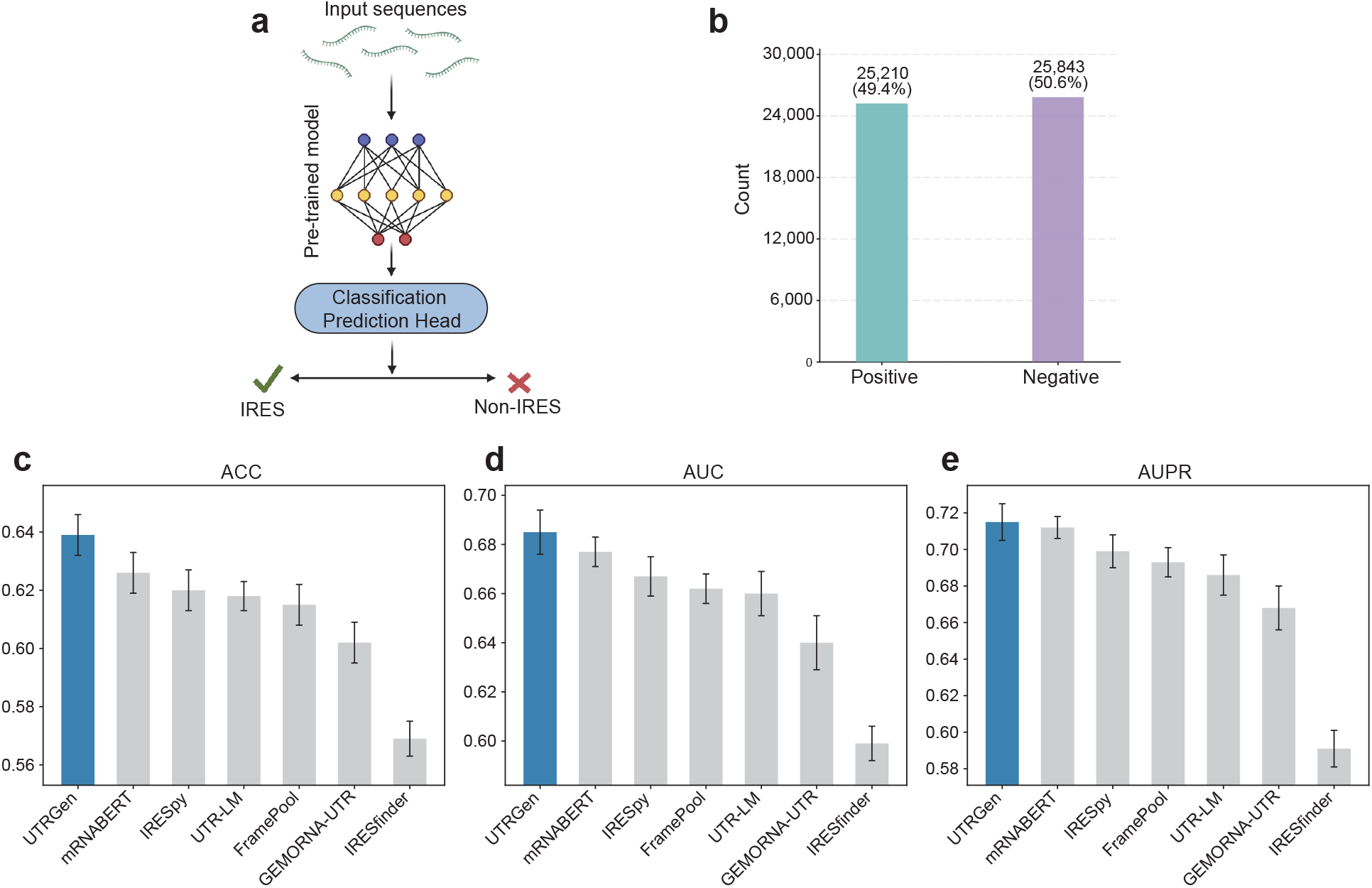
UTRGen enables accurate identification of IRESs. **a**, Task setup for IRES prediction by fine-tuning the pre-trained UTRGen with a classification head. **b**, Composition of the IRES benchmark dataset (25,210 positives and 25,843 negatives). **c**–**e**, Ten-fold cross-validation performance comparison of UTRGen with six baselines evaluated by ACC (**c**), AUC (**d**), and AUPR (**e**). Bars represent the mean across folds, and error bars indicate the s.d. Paired *t*-tests on fold-wise results supported significant improvements for UTRGen in all ACC and AUC comparisons and in five of six AUPR comparisons (*P <* 0.05); the only non-significant AUPR comparison was against mRNABERT. Icons in figures were created with BioRender.com.

For benchmarking, we compared UTRGen with six baseline methods, including three pre-trained mRNA language models (mRNABERT [11], UTR-LM [10], and GEMORNA-UTR [12]), two task-specific predictors (IRESpy [27] and IRESfinder [28]), and the feature-pooling method FramePool [8], all evaluated under the same ten-fold cross-validation protocol to ensure consistency. As shown in Fig. 5c–e and Supplementary Table 5, UTRGen achieved the best performance across all evaluation metrics, with an accuracy (ACC) of 0.639, an area under the receiver operating characteristic curve (AUC) of 0.685, and an area under the precision–recall curve (AUPR) of 0.715. Compared with the strongest baseline, mRNABERT [11], UTRGen improved ACC by 2.1%, AUC by 1.2%, and AUPR by 0.4%. Larger gains were observed relative to other baselines; for example, compared with IRESfinder [28], UTRGen improved ACC by 12.3%, AUC by 14.4%, and AUPR by 21.0%. Paired *t*-tests on fold-wise cross-validation results showed that UTRGen significantly outperformed all six baselines in ACC and AUC (*P <* 0.05 for all comparisons), and significantly outperformed five of six baselines in AUPR, with the only non-significant difference observed against mRNABERT (*P* = 0.4289; Supplementary Table 6). These results indicate that UTRGen more effectively captures sequence features associated with IRES activity than existing methods.

### 2.6 Enabling constraint-aware and function-guided 5′UTR design via reinforcement learning

Among the downstream properties modeled by UTRGen, we focused on TE and EL as function-guided design targets because they provide biologically meaningful and application-relevant readouts of 5′ UTR activity. TE was prioritized as the primary optimization objective because it directly reflects translational output and is therefore closely aligned with the central goal of improving protein production in mRNA engineering. EL was included as a complementary expression-related objective to assess whether the RL framework could generalize beyond TE-oriented optimization within the same cellular context.

We selected the human embryonic kidney-derived HEK293T cell line as the primary testbed for RL-driven, function-guided 5′ UTR design for three reasons. First, HEK293T is a widely used and well-characterized mammalian transfection system and is commonly used in reporter assays and studies of translational regulation. Second, HEK293T provides a standardized and practically relevant starting point for mammalian expression optimization, as well as a useful reference for testing generalization to other cell types or conditions. Third, this choice follows common mRNA engineering practice in which rapid iteration is first performed in a standardized system such as HEK cells and then extended to more application-relevant contexts.

#### RL optimization under multi-objective constraints converges stably and improves TE consistently

In the HEK293T setting, we treated TE as the primary optimization objective and used UTRGen-TE (HEK) as the reward model. To restrict the search to a more realistic and biophysically feasible sequence/structure regime, we explicitly incorporated constraints on GC content (0.4–0.6) and MFE into the reward function. The GC constraint was designed to suppress systematic compositional drift and prevent the model from deviating from typical nucleotide composition ranges. The MFE constraint was introduced to limit excessive secondary-structure stability in the 5′ UTR, thereby reducing the likelihood that the agent would obtain spuriously high scores by exploiting extreme structural patterns. The RL training trajectory (Fig. 6a) shows stable convergence under these multi-objective constraints: both the total weighted reward and the predicted TE increase steadily, while GC content and MFE remain stable throughout optimization, indicating that the performance gains are not achieved at the expense of basic compositional or structural plausibility.

**Figure 6.**
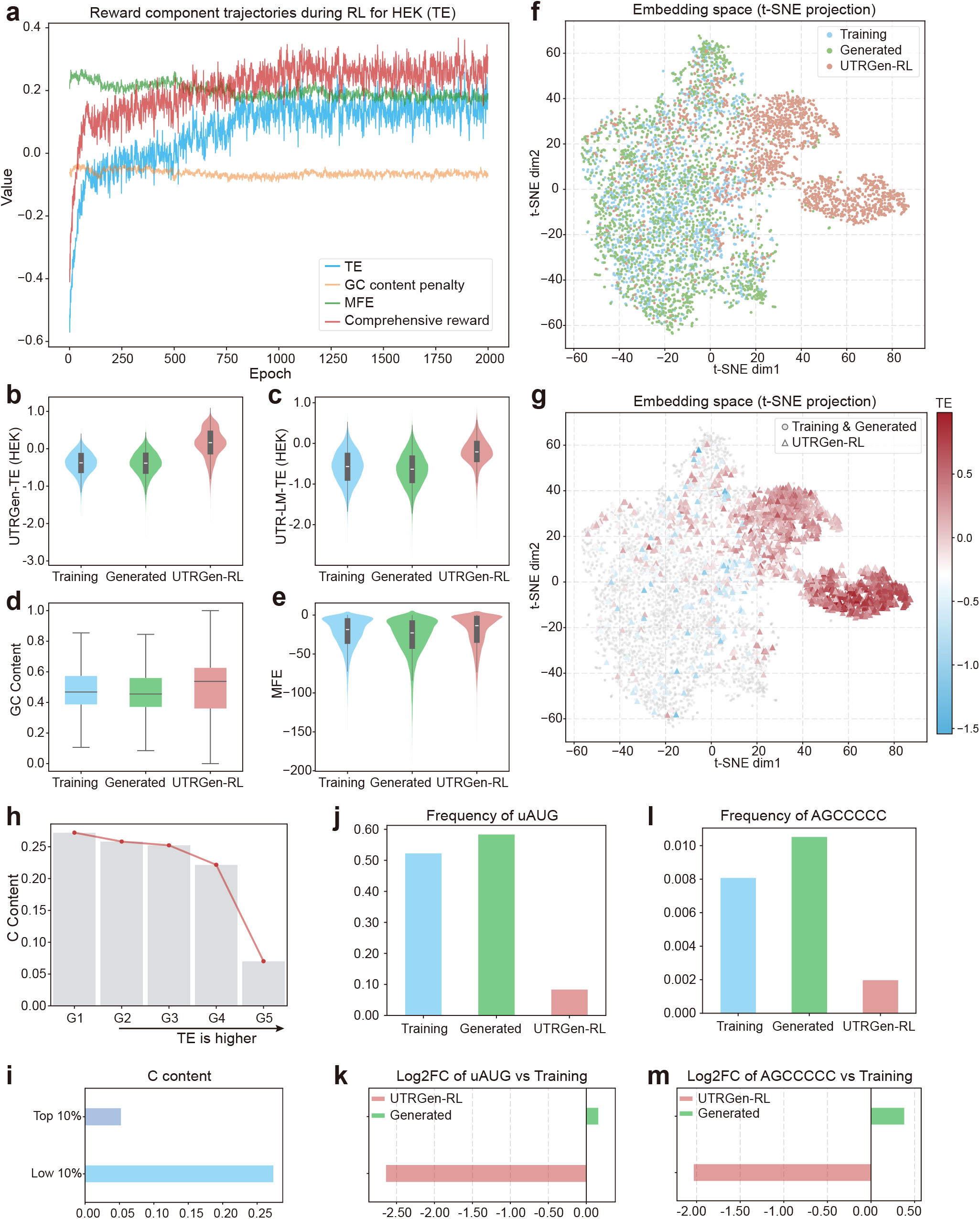
KL-regularized RL enables constraint-aware, TE-guided 5′ UTR design. **a**, Trajectories of the reward components during RL training, including the predicted TE reward, GC-content penalty, MFE-related term, and the total weighted reward, showing stable convergence under multi-objective constraints. **b, c**, Distributions of predicted TE scores for Training, Generated and RL-optimized sequences (UTRGen-RL), evaluated using the internal reward model UTRGen-TE (HEK) (**b**) and an independent predictor UTR-LM-TE (HEK) (**c**). **d, e**, GC content (**d**) and MFE (**e**) distributions across Training, Generated, and UTRGen-RL, showing that RL improves TE while maintaining native-like composition and secondary-structure stability. **f**, t-SNE projection of final-layer sequence embeddings for Training, Generated, and UTRGen-RL sequences, highlighting a distributional shift of RL-optimized sequences toward a distinct region in representation space. **g**, The same embedding space coloured by predicted TE, showing that the RL-enriched region coincides with high predicted TE. **h, i**, Cytidine content decreases with increasing predicted TE across UTRGen-RL deciles (**h**), and the top 10% UTRGen-RL sequences show lower C content than the bottom 10% (**i**). **j, k**, Frequency of uAUG across sets (**j**) and depletion quantified by log2FC relative to Training (**k**). **l, m**, Frequency of the inhibitory motif AGCCCCC across sets (**l**) and depletion quantified by log2FC relative to Training (**m**).

To further quantify the optimization effect and mitigate concerns about reward hacking or overfitting to the internal scorer, we compared score distributions across training sequences (Training), UTRGen-generated sequences (Generated), and RL-optimized sequences (UTRGen-RL) using multiple TE prediction tools. In addition to evaluation with the reward model UTRGen-TE (HEK) (Fig. 6b), we performed orthogonal assessments using three independent third-party predictors, UTR-LM-TE (HEK), mRNABERT-TE (HEK), and GEMORNA-TE (HEK) (Fig. 6c and Supplementary Fig. 3a, b). All of these predictors consistently show an upward shift in the TE distribution for UTRGen-RL relative to both Training and Generated sequences. The agreement across predictors with different architectures and training priors provides convergent computational support for the robustness of the optimization outcome. Meanwhile, the distributions of GC content (Fig. 6d) and MFE (Fig. 6e) for UTRGen-RL remain on the same scale as those of the training set, supporting the conclusion that the TE improvement stems from goal-directed search rather than degenerate, constraint-violating sequences.

To further dissect the contribution of individual reward components to the optimization behavior, we performed reward-function ablation experiments by removing the GC constraint, the MFE constraint, or both constraints while retaining only the TE reward for training and evaluation. The results showed that optimizing TE alone yielded the highest target scores, but produced sequences with GC-content distributions that deviated markedly from those of natural 5′ UTRs. Adding either constraint partially corrected the compositional or structural properties of the generated sequences, although noticeable deviations still remained. In contrast, the full reward formulation, which jointly incorporated both GC and MFE constraints, preserved relatively high TE prediction scores while producing sequences whose GC-content and MFE distributions were more consistent with those of natural 5′ UTRs, indicating that this design achieves a better balance between target optimization and sequence plausibility (see Supplementary Note 1.3.2 and Supplementary Fig. 4).

#### RL explores the representation landscape and enriches a high-fitness cluster associated with predicted TE

To visualize how RL alters the distribution of sequence representations, we extracted embeddings from the final layer of the model for sequences of Training, Generated, and UTRGen-RL, followed by t-SNE projection and clustering (see Supplementary Methods). As shown in Fig. 6f, generated sequences (Generated, green) largely overlap with natural training sequences (Training, blue) in the latent space, consistent with the pre-trained generator primarily sampling along the empirical data manifold. In contrast, RL-optimized sequences (UTRGen-RL, red) form a more compact cluster outside the original Training/Generated distribution, indicating that RL drives the generative distribution toward a distinct region of representation space. When colored by predicted TE from the reward model UTRGen-TE (HEK) (Fig. 6g), this emergent cluster is enriched for high-scoring sequences. Importantly, the same spatial pattern is recapitulated when the embedding landscape is colored using three independent TE predictors, UTR-LM-TE (HEK), mRNABERT-TE (HEK), and GEMORNA-TE (HEK) (Supplementary Fig. 3c–e), indicating that the RL-enriched cluster is consistently associated with elevated predicted TE across models. Together, these results support the interpretation that, under multi-objective constraints, RL identifies and enriches a distinct “high-fitness cluster” associated with the target phenotype of elevated TE, rather than a purely model-specific optimum.

#### RL generalizes beyond TE and enables stable EL-guided 5′UTR design

In addition, to examine whether the RL framework can generalize to different optimization objectives within the same cellular context, we also performed RL-based 5′ UTR design in HEK293T cells using EL as the target. The results showed trends consistent with those observed in the TE optimization setting: under multi-objective constraints, the model exhibited stable training dynamics and shifted the optimized sequence distribution toward regions with higher predicted EL scores (Supplementary Fig. 5). These findings suggest that UTRGen-RL is not limited to TE-oriented design, but can also be adapted to other expression-related optimization objectives.

#### Reproducible trends in a more specialized, tumor-relevant context (PC3)

Beyond HEK293T, we performed a corresponding set of experiments in a more specialized prostate cancer–relevant cellular background, the human prostate cancer cell line PC3, as described in Supplementary Fig. 6. Overall, the PC3 setting recapitulates the major trends observed in HEK293T. RL training remains stable under the same constraints. RL-optimized sequences show a global increase in TE-related scores (Supplementary Fig. 6a–c). Summary statistics for constraints such as GC content and MFE do not exhibit systematic drift (Supplementary Fig. 6d, e). Latent-space analysis further indicates a similarly pronounced distributional shift of RL-optimized sequences relative to generated sequences, with higher-density enrichment in regions associated with high predicted TE (Supplementary Fig. 6f, g). Collectively, these results support that our framework shows consistent goal-directed optimization across cellular contexts and demonstrates a degree of transferability.

### 2.7 Interpreting key translation-associated features and motifs in RL-designed 5′UTRs

To improve the interpretability of how RL achieves a higher TE, we systematically analyzed sequence-level features and regulatory motifs in the UTRGen-RL designs. In the HEK293T setting, UTRGen-RL sequences showed an overall decrease in cytidine content. When we divided the UTRGen-RL sequences into five equal groups based on their predicted TE (ranging from low to high), the average C content within each group exhibited a negative correlation with TE (Fig. 6h). The top 10% sequences had significantly lower C content than the bottom 10% sequences (Fig. 6i). This preference, which emerged during goal-directed optimization, is consistent with previous reports that higher C content is associated with reduced ribosome recruitment and lower translation [15]. These results suggest that the model captures compositional constraints relevant to translational regulation without explicitly encoding such rules. A consistent trend in C content was also observed in the PC3 setting (Supplementary Fig. 7a, b).

We next used log2 fold change, log2FC, to quantify enrichment and depletion of key translation-associated motifs before and after RL optimization, comparing Generated and UTRGen-RL sequences. Based on the canonical scanning model of translation initiation, uAUGs can act as premature start codons that compete for scanning ribosomes, thereby reducing initiation at the main open reading frame and suppressing coding-sequence translation [16]. Consistent with this expectation, uAUG frequency was markedly reduced in UTRGen-RL sequences (Fig. 6j) and was strongly depleted after RL (log2FC = -2.64; Fig. 6k), indicating that RL tends to avoid unfavorable initiation signals. The same pattern was observed for PC3-optimized sequences (Supplementary Fig. 7c, d).

Beyond uAUGs, we examined previously reported inhibitory motifs. A multi-omics analysis by Eraslan *et al*. reported that the motif AGCCCCC is associated with translational repression in kidney cells [17]. In line with this, RL optimization targeting HEK293T TE led to significant depletion of AGCCCCC (log2FC = -2.03; Fig. 6l, m), suggesting that the learned strategy reduces sequence patterns linked to translational inhibition. In the PC3 setting, the same study further reported that AGCGGAA and AGCAAC are associated with translational repression in prostate cells [17]. Correspondingly, RL optimization targeting PC3 TE depleted these motifs, with log2FC values of -2.82 and -0.89, respectively (Supplementary Fig. 7e–h). Overall, UTRGen-RL shows directional changes in nucleotide composition and motif usage that are consistent with established principles of translational control, providing interpretable sequence-level support for the RL optimization outcomes.

## 3 Discussion

In this study, we developed UTRGen, a unified framework for full-spectrum mRNA 5′ UTR design that integrates sequence generation, multi-property prediction, and function-guided design within a single system. Previous computational approaches have largely focused on passive functional prediction or heuristic optimization within limited natural sequence libraries. By contrast, UTRGen enables active, goal-directed exploration of a substantially broader 5′ UTR sequence space, thereby extending computational design from sequence-function inference to controllable *de novo* design.

The generative capability of UTRGen is supported by autoregressive pre-training on more than 9.4 million eukaryotic 5′ UTR sequences from multiple species. Our analyses indicate that the model captures conserved and transferable regulatory features rather than merely reproducing training-set sequences. In particular, UTRGen preserves both local motif preferences and higher-order sequence properties, including GC-content distribution and thermodynamic stability. As a result, it can generate novel and diverse candidate sequences that remain consistent with natural 5′ UTRs in their distributional, structural, and functional characteristics, supporting their use as biologically plausible designs.

On the basis of this pre-trained representation, UTRGen also achieves strong performance across diverse downstream prediction tasks, including MRL, TE, EL, and IRES activity. Across these benchmarks, the framework consistently performs at or above existing baselines, with especially strong gains in TE and EL prediction. These results suggest that the learned representation captures sequence determinants relevant to multiple aspects of translational regulation. Beyond predictive accuracy alone, this capability is important because it provides a reliable reward signal for subsequent function-guided design.

To extend UTRGen beyond functional prediction toward function-guided sequence design, we incorporated GRPO-based reinforcement learning into the framework. This module enables the model to generate sequences with improved TE and EL while preserving key constraints on sequence composition and structural properties. We selected TE and EL as optimization targets because they capture two complementary and practically important aspects of 5′ UTR function: TE more directly reflects translational regulation, whereas EL reflects the overall expression output that is often more relevant for downstream applications. The stable optimization trajectories observed during training, together with the emergence of clusters enriched for high-TE/EL sequences in latent space, suggest that UTRGen-RL can identify favorable regulatory patterns that are rarely sampled in natural sequence space. Importantly, the optimized sequences do not appear to arise from trivial reward exploitation. Instead, they remain consistent with known principles of translational regulation, including depletion of C content, uAUGs, and inhibitory motifs, supporting the biological interpretability of the learned design strategy.

These results highlight the potential of UTRGen for therapeutic mRNA engineering and related applications in synthetic biology. The ability to generate non-natural 5′ UTRs that satisfy both functional objectives and biological constraints may be particularly valuable for improving translational output in a controlled manner. In addition, the transferability of the framework across distinct cellular contexts, including HEK293T and PC3 cells, suggests its potential utility for context-specific or cell-type-aware mRNA design.

Overall, we developed UTRGen as a unified and extensible computational framework for full-spectrum 5′ UTR design, integrating sequence generation, multi-property prediction, and function-guided design within a single architecture. Our results show that UTRGen can generate novel, diverse, and biologically plausible 5′ UTRs, accurately predict multiple translation-related properties, and guide the design of sequences with desirable functional characteristics. More broadly, this work advances 5′ UTR engineering from fragmented sequence profiling and candidate screening toward unified computational design, and provides a potentially generalizable strategy for the programmable design of other cis-regulatory elements, with broader implications for sequence design across RNA, DNA, and protein engineering.

## 4 Methods

### 4.1 Datasets

#### pre-training datasets

To ensure UTRGen adequately captures the underlying sequence syntax and latent features of mRNA 5′ UTRs via autoregressive pre-training, we assembled a large corpus of 5′ sequences from two public resources. The first, UTRdb [18], is a dedicated database of untranslated regions, cataloging an extensive array of 5′ and 3′ UTR sequences from 573 eukaryotic species. And second, the NCBI [19], a foundational database resource for biomedicine and genomics. After extraction, the 5′ UTR sequences underwent rigorous preprocessing, including quality control, redundancy removal, and sequence clustering. This curation process generated a refined corpus of 9,446,225 unique 5′ UTR sequences. For the pre-training phase, we partitioned this corpus into a training set of 9,361,120 sequences and a test set of 85,105 sequences. Details of data collection and preprocessing are provided in Supplementary Note 1.1.1.

#### Downstream datasets

For MRL prediction, we employed eight fixed-length synthetic 5′ UTR libraries (U_1_, U_2_, Ψ_1_, Ψ_2_, mC-U_1_, mC-U_2_, m1Ψ_1_, and m1Ψ_2_). Each library comprises approximately 280,000 random sequences of length 50 nt [5]. To further evaluate performance on variable-length sequences, we incorporated two datasets originally proposed by Sample *et al*. [5]. These collections consist of 83,919 random 5′ UTR sequences and 15,555 human 5′ UTR sequences, spanning lengths from 25 to 100 nt. Additional details are provided in Supplementary Table 7.

For TE and EL prediction, we used three endogenous human 5′ UTR datasets derived from distinct cell line or tissue type [6]: human muscle tissue (muscle), the human prostate cancer cell line PC3, and the human embryonic kidney (HEK) 293T cell line. Dataset details are provided in Supplementary Table 8.

For the IRES prediction task, we constructed a binary sequence dataset by integrating records from multiple sources, including IRESbase [32], Rfam [33], IRESite [34], IRESPred [31], Weingarten-Gabbay *et al*. [35], and Chen *et al*. [36]. All collected sequences were subjected to unified filtering and deduplication to remove redundant entries across sources while preserving a relatively balanced class distribution. The final dataset contained 51,053 sequences, including 25,210 IRES-positive and 25,843 non-IRES-negative sequences. The average sequence length was approximately 190 nt for IRES sequences and 179 nt for non-IRES sequences. Further details on sequence collection and preprocessing are provided in Supplementary Note 1.1.2.

### 4.2 Architecture and pre-training of UTRGen

The proposed UTRGen employs a decoder-only Transformer architecture based on the GPT-2 framework. As illustrated in Fig. 1a, the pre-training model comprises eight stacked masked self-attention blocks followed by a linear output head. The model uses 16 attention heads, a hidden size of 384, and a maximum sequence length of 256, for a total of 14.3 million trainable parameters. To represent nucleotide sequences, we adopted the tokenization scheme described by Chu *et al*. [10]. The vocabulary *V* comprises seven tokens: four canonical nucleotides ({ *A,U,C, G* }) and three special tokens ([PAD], [BOS], [EOS]). During preprocessing, a [BOS] token is prepended to denote the start of the sequence, and an [EOS] token is appended to signify its termination. Inputs are truncated to a maximum of 256 nt, whereas shorter sequences are padded to this length using [PAD] tokens. Tokenized sequences were mapped to continuous high-dimensional embeddings and combined element-wise with positional encodings. The resulting vectors then propagate through the decoder layers, with the final hidden states routed to an output layer for autoregressive next-token prediction (NTP) during pre-training.

#### Next token prediction

NTP serves as the primary pre-training objective for UTRGen, formulated to predict the subsequent nucleotide within a 5′ UTR sequence given its preceding context. Specifically, given a partial 5′ UTR sequence *X* = {*x*_1_, *x*_2_, …, *x*_*t*−1_}, the model infers the identity of the next token *x*_*t*_ ∈ {*A,U,C, G*, [EOS]}. This objective enables the model to capture sequential dependencies and contextual patterns in 5′ UTRs. The training objective was defined using the cross-entropy loss. For a given sequence *X* = {*x*_1_, *x*_2_, …, *x*_*T*_ }, the loss is defined as:

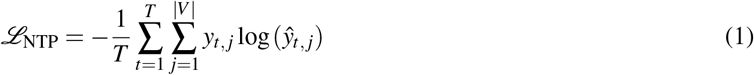

where *T* represents the total sequence length, and | *V* | defines the vocabulary size. Here, *y*_*t, j*_ is a one-hot indicator of the ground-truth token at position *t* (evaluating to 1 for the correct target and 0 otherwise), and *ŷ*_*t, j*_ is the predicted probability of token *j* at that position. Crucially, indices corresponding to [PAD] tokens are explicitly masked during this computation, ensuring they are excluded from the gradient backward propagation.

### 4.3 Downstream Tasks

After obtaining the pre-trained embedding model, we further explore its applicability in various downstream tasks related to 5′ UTR, mainly including 5′ UTR generation and function prediction (Fig. 1a, b).

#### 5′ UTR generation

For the sequence generation task, we assessed whether UTRGen could generate novel and diverse 5′ UTR sequences while preserving the structural and functional characteristics of natural 5′ UTRs. To examine performance across different sequence spans, we generated sequences in four length intervals: 25-50 nt, 50-100 nt, 100-150 nt, and 150-200 nt. Generation was initiated from a single randomly selected nucleotide and proceeded autoregressively. For each interval, 10,000 unique sequences were sampled, resulting in 40,000 *de novo*-generated sequences in total. For comparison, we constructed length-matched natural reference datasets by stratified random sampling from the pre-training training and test sets. For each length interval, 10,000 5′ UTR sequences were sampled from each split, yielding two reference sets of 40,000 sequences each. These datasets were used to evaluate novelty, diversity, and similarity to natural sequences in structural and functional properties. Further details are provided in Supplementary Note 1.2.

#### 5′ UTR function prediction

Given the pivotal role of the 5′ UTR in translational regulation, we initially probed UTRGen’s utility across four key functional prediction tasks: MRL, TE, EL, and IRES recognition. Specifically, MRL quantifies the concurrent ribosomal load on a given mRNA; TE estimates the overall efficiency of protein synthesis; EL profiles the relative intracellular transcript abundance; and IRES elements drive cap-independent ribosomal recruitment. Collectively, these metrics provide a holistic functional characterization of 5′ UTR sequences.

To rigorously test the representational power of the embeddings learned during pre-training, we deliberately implemented a simple predictor design for downstream fine-tuning. High-dimensional semantic features from the terminal decoder layer were channeled into a straightforward two-layer 1D convolutional neural network followed by a linear output layer. The model produced scalar outputs for the regression tasks (MRL, TE, and EL) and class probabilities for the classification task (IRES prediction).

### 4.4 Reinforcement learning for function-guided 5′ **UTR design**

We implemented an RL fine-tuning framework based on GRPO [13, 14] to steer the pre-trained UTRGen model to generate *de novo* sequences with optimized functional properties, particularly enhanced TE and EL. The autoregressive generation of 5′ UTRs was formulated as a Markov decision process (MDP). Within this RL paradigm, given a generated nucleotide prefix as the current state *s*_*t*_, and the policy model *π*_*θ*_, initialized from the pre-trained UTRGen weights, selects the next nucleotide *a*_*t*_ from the discrete action space {*A,U,C, G*}. At the start of each generative episode (*t* = 0), the initial state consisted of a single randomly sampled nucleotide used as a prompt. Because the biological efficacy of an mRNA transcript can only be evaluated from the complete sequence, intermediate rewards were set to 0, and the terminal reward *R*_*T*_ was computed strictly upon generation completion.

We adopted the light GRPO algorithm to avoid the substantial GPU memory overhead associated with the value network in conventional PPO, which often has a parameter count comparable to that of the policy model itself. At each optimization step, the model generated a group of candidate sequences (*G*) independently from the same single-nucleotide prompt. We leverage the intra-group reward mean *µ*_group_ of these *G* sequences to establish a dynamic baseline. Because biological evaluation metrics for 5′ UTR generation exhibit substantial variance and can change sharply in response to single-nucleotide mutations, the advantage scores were further normalized using the reward standard deviation across the full training batch, *σ*_batch_, to improve training stability [37]. The advantage score for the *i*-th sequence, *A*_*i*_, was therefore defined as:

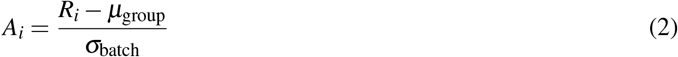

For parameter updates, we used an on-policy gradient formulation to maximize the expected advantage *A*_*i*_. To prevent the policy from deviating excessively from native biological sequence syntax while pursuing higher rewards, the objective included a KL divergence penalty that constrained the active policy against a frozen pre-trained reference model, *π*_ref_. The final optimization objective was defined as:

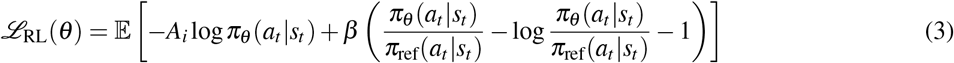

where *β* is a scaling hyperparameter controlling the strength of the KL penalty. Details of its tuning are provided in Supplementary Note 1.3.1.

#### Multi-objective reward function

We defined a dynamically adjustable multi-objective reward function to comprehensively evaluate generated sequences. The reward comprised three components: a target functional reward (*r*_Func_), GC content penalty (*r*_GC_Penalty_), and a thermodynamic stability reward (*r*_MFE_). To compute the target functional reward, we used the task-specific UTRGen-TE or UTRGen-EL model, with weights frozen, to perform forward inference on each generated sequence. The resulting continuous scalar output was used directly as the reward score:

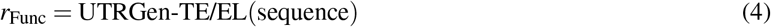

To constrain compositional bias, we defined an optimal GC-content interval of [0.4, 0.6] and imposed a penalty on sequences falling outside this range:

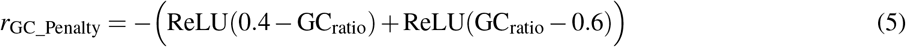

Thermodynamic stability was evaluated using the ViennaRNA package [38] to calculate the MFE. To reduce sequence-length effects, the MFE was normalized by sequence length:

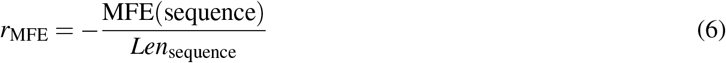

The global reward was defined as a weighted linear combination of these components,

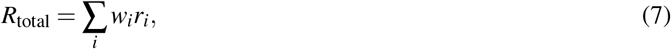

where *r*_*i*_ denotes one of the individual reward terms, including *r*_Func_, *r*_GC_ and *r*_MFE_, and *w*_*i*_ denotes the corresponding weighting coefficient. By modulating the weighting coefficients, the optimization trajectory could be flexibly adjusted to steer sequence-space exploration toward specific biological phenotypes. Details of the *w*_*i*_ settings are provided in Supplementary Note 1.3.3.

### 4.5 Experimental setting

UTRGen was trained in three stages: autoregressive pre-training, downstream task-specific fine-tuning, and reinforcement learning optimization. Pre-training was performed with the AdamW optimizer and cosine annealing on two NVIDIA A100 GPUs for 100 epochs. For downstream function prediction, the pre-trained model was fine-tuned on individual tasks using AdamW-based optimization, with regression tasks trained using the Huber loss; downstream training and evaluation were conducted on a single NVIDIA A100 or A40 GPU. Reinforcement learning optimization was performed using GRPO on a single NVIDIA A100 GPU. Experimental details and hyperparameters are provided in Supplementary Note 1.3 and 1.4.

## Data availability

The datasets used in this study are available at https://drive.google.com/drive/folders/1E4TYXN48wdDru_-ETVxu2UWYSEKvO0wr?usp=sharing, which includes the pre-training dataset and datasets for the downstream tasks. Detailed dataset statistics are provided in Supplementary Note 1.1. Source data are provided with this paper.

## Code availability

The latest version of the code is available at https://github.com/HIM-AIM/UTRGen under the MIT license. The pre-trained model checkpoint is available at https://drive.google.com/drive/folders/1E4TYXN48wdDru_-ETVxu2UWYSEKvO0wr?usp=sharing.

## Acknowledgements

This work was supported by the Zhejiang Provincial Science and Technology Program (2025C01129 to X.L.), the China Postdoctoral Science Foundation (2025M772773 to Z.W.), the National Natural Science Foundation of China (32401159, 32571597 to J.H.), and the Zhejiang Provincial Natural Science Foundation of China (QKHM25B0502 to J.H.).

## Author contributions

Z.W., X.L., and J. H. conceptualized the study, designed and developed the UTRGen model, and conducted the investigation; Z.W., M.C., and X.Z. curated data, performed the evaluation, analyzed the data, and wrote the code; Z.W. and M.C. wrote the initial draft of the manuscript; X.L., J.H., and J.Z. supervised the overall study; and all authors reviewed and revised the manuscript.

## Supplementary Information

### 1 Supplementary Note

#### 1.1 Data Description

##### 1.1.1 Details on pre-training data collection and processing

This section details the comprehensive pipeline used to construct the large-scale, high-quality eukaryotic 5′ UTR dataset for UTRGen’s autoregressive pre-training.

###### Data Acquisition and Initial Compilation

Raw sequence data were gathered from two authoritative nucleic acid databases to maximize species diversity and annotation reliability:

- UTRdb [18] is a comprehensive database specialized for UTR sequences, providing rich annotations such as regulatory elements, conserved regions, and experimentally validated miRNA targets. We retrieved data files for all 573 species, which include 5′ UTR, 3′ UTR, and CDS sequences for each transcript. From these records, we extracted approximately 18 million 5′ UTR sequences. By merging sequences based on transcript and gene identifiers (rejoining segments originally separated by introns), we obtained roughly 12 million complete 5′ UTR sequences.
- We augmented our dataset from the NCBI database [19] with complete mRNA transcripts from five key model organisms (human, mouse, rat, chicken, and zebrafish). Utilizing genomic annotations, we localized the transcription start sites (TSS) and start codons (AUG) to precisely cut out the 5′ UTR regions, contributing an additional 577,694 sequences. Merging these sources established an initial data pool of 13,153,185 raw sequences.

###### Quality Control and Redundancy Filtration

To ensure corpus quality, we applied a rigorous filtering pipeline to the initial sequence pool. Sequences were excluded if they lacked complete annotations, contained ambiguous nucleotides (for example, N), or were shorter than 15 nt. After filtering, 12,413,620 high-quality sequences were retained.

###### Homology-Based Sequence Clustering

To reduce distribution bias and the overrepresentation of homologous elements, the filtered sequences were clustered using MMseqs2 [39]. Parameters were configured for a minimum sequence identity of ≥ 80% and a coverage threshold of ≥ 80% (cluster mode = 2). This step produced 8,510,886 independent clusters. Within each cluster, the longest sequence was designated as the representative element.

###### Dataset Splitting and Stratified Sampling

For the final pre-training corpus construction, all clusters were randomly partitioned into training (99%, yielding 8,425,777 clusters) and validation (1%, yielding 85,105 clusters) pools. We then executed the following stratified sampling strategy:

- Training Set: The representative sequence from each training cluster was included. To preserve a reasonable range of natural variation, we randomly sampled 50% of the remaining intra-cluster members (capped at a maximum of 200 sequences per cluster) and incorporated them into the training set.
- Test Set: Only the representative sequences from the validation clusters were strictly retained. This rigorous separation guarantees absolute sequence non-redundancy between the training and validation sets, ensuring an unbiased evaluation of the model’s generalization capabilities.

Through this workflow, we finalized a comprehensive dataset of 9,446,225 unique natural 5′ UTR sequences, comprising 9,361,120 training and 85,105 test sequences. This corpus provides the foundational diversity necessary for UTRGen to accurately model natural sequence distributions.

#### 1.1.2 Details of the IRES dataset collection and processing procedures

This section describes the IRES dataset we have compiled. To systematically evaluate the model’s generalization ability in the IRES classification task, we integrated multiple public databases and published experimental data to construct a large-scale, deduplicated, and bias-controlled IRES classification dataset. Specifically, these data are sourced from IRESbase [32], Rfam [33], IRESite [34], IRESPred [31], Weingarten-Gabbay *et al*. [35], and Chen *et al*. [36]. During the initial data screening phase, we extracted sequences explicitly annotated as having IRES activity from the above sources to form a positive candidate set, while the remaining sequences without IRES functional annotations were classified as a negative candidate set.

For the initially compiled sequence set, we first performed a rigorous label conflict check to remove anomalous sequences that appeared in both the positive and negative sets. Subsequently, a length filter (6-1500 nt) was applied to all sequences. This range sufficiently covers the length distribution of the vast majority of known eukaryotic and viral IRESs and effectively avoids the negative impact of extremely long sequences on model training stability. To eliminate data bias caused by homologous sequences, we performed clustering and deduplication using the MMseqs2[39] tool (parameter settings: cov-mode=1, cluster-mode=2, sequence identity ≥0.5).

Following the above standardization process, the final large-scale IRES dataset comprised 51,053 high-quality sequences. Specifically, the positive sample set comprises 25,210 IRES sequences with an average length of 189.5 nt, while the negative sample set contains 25,843 sequences with an average length of 178.9 nt. Detailed distributions of sequence lengths and sources are presented in Supplementary Table 9.

### 1.2 Details on the Sequences Used to Evaluate Sequence Modeling Capabilities

To comprehensively and unbiasedly evaluate UTRGen’s ability to capture natural 5’ UTR sequence syntax and its generative capacity, we constructed a structured comparative analysis dataset across different length distributions. This dataset consists of two parts: synthetic sequences generated de novo by the model, and natural reference sequences used as a baseline.

#### *De novo* generated sequences

To explore the model’s performance across different sequence spans during the generation phase, we divided our generation targets into four specific length intervals: 25-50 nt, 50-100 nt, 100-150 nt, and 150-200 nt. We used a single random nucleotide (A, U, C, or G) as the initial prompt during the inference phase. This guided the model’s unconditional autoregressive generation. Ultimately, we independently sampled 10,000 completely unique sequences within each of the four length intervals, producing a total of 40,000 generated sequence samples.

#### Natural reference sequences

To conduct rigorous comparisons of physical properties and sequence syntax with the generated sequences, we constructed a natural sequence baseline set of matching scale. We performed random stratified sampling without replacement from both the training and test sets used in the pre-training phase. Matching the settings of the generated sequences, we randomly sampled 10,000 real 5′ UTR sequences from each of the same four length intervals (25-50 nt, 50-100 nt, 100-150 nt, and 150-200 nt). This reference dataset contains a total of two samples, each consisting of 40,000 natural sequences.

Through this length-matched sampling approach, we compiled a total of 120,000 high-quality sequences for analysis. This provides reliable data to support the subsequent evaluation of the generated sequences’ novelty, diversity, and fidelity.

### 1.3 Ablation Experiments on Reinforcement Learning Training Parameters

To systematically validate the necessity of multi-objective constraints and the robustness of hyperparameter selection within the RL framework, we conducted ablation studies and boundary analyses using the HEK293T cell line. This analysis aimed to ensure that, while maximizing the target TE, the generative model strictly adheres to the biophysical distribution characteristics of natural 5′ UTRs, thereby avoiding the generation of biologically unfeasible sequences.

#### 1.3.1 Sensitivity Analysis of the KL Divergence Penalty

We first conducted a sensitivity analysis on the KL divergence penalty coefficient (*β*) within the GRPO algorithm. This coefficient directly dictates the extent to which the generative policy adheres to the prior distribution of pre-trained natural sequences. As illustrated in Supplementary Fig. 4a, a pronounced trade-off exists between goal-oriented optimization and sequence diversity. When employing a lower penalty coefficient (*β* = 0.3), the model exhibits an aggressive tendency toward TE growth (mean TE = 0.49). However, this comes at the cost of a sharp decline in the average Shannon entropy of the generated sequences to 0.82. This indicates severely compromised sequence diversity, suggesting the model has fallen into mode collapse by generating simple, repetitive sequences. Conversely, an excessively high coefficient (*β* = 0.7) improves the entropy (mean entropy = 1.29) but still falls short of the natural sequence entropy level (1.89). At the same time, this overly stringent KL constraint severely impairs the agent’s ability to explore novel high-TE sequence spaces, leading to a drastically diminished functional gain (mean TE of only 0.10). Balancing these considerations, we established *β* = 0.4 as the final benchmark parameter. This setting achieves a substantial and stable functional enhancement in TE while preserving relatively high sequence diversity. Consequently, it maintains the requisite exploration space for subsequent model generation constrained by the composite reward function.

#### 1.3.2 Ablation of Reward Components

After establishing the KL penalty baseline, we conducted ablation studies on the core components of the reward function to validate the necessity of multi-objective constraints within the reinforcement learning framework (Supplementary Fig. 4 b-d). When relying solely on predicted TE as the sole reward signal (TE-Only), the model generated extremely anomalous sequences in an attempt to achieve high scores. These sequences not only exhibited abnormally low GC content, averaging 0.26 (Supplementary Fig. 4b), but also nearly lost all secondary structure, with an average minimum free energy (MFE) of only -3.03 kcal/mol (Supplementary Fig. 4c), completely deviating from the physical characteristics of natural 5′ UTRs. Furthermore, applying individual physical constraints also revealed severe limitations. When only the GC penalty was applied (TE+GC), the sequence structures remained overly unstructured (Supplementary Fig. 4c). Conversely, when only the MFE constraint was applied (TE+MFE), the model excessively stacked G-C base pairs to maximize the thermodynamic stability score, driving the GC content up to a non-physiological state above 0.75 (Supplementary Fig. 4b). Only under the complete, synergistic reward configuration (TE + GC penalty + MFE constraint) were these “reward hacking” pathways entirely blocked. While achieving substantial TE gains (Supplementary Fig. 4d), the sequences generated under this full configuration maintained a mean GC content (0.50) and a mean MFE (-20.77 kcal/mol) that perfectly aligned with the training set distribution of native natural sequences (Supplementary Fig. 4b–c).

#### 1.3.3 Reward Weight Analysis

Finally, to determine the balance between functional objectives and structural constraints, we conducted an objective boundary analysis of the weights for the GC penalty and the MFE reward (Supplementary Table 10). For the GC constraint, we compared different levels of GC penalty while keeping the MFE reward weight fixed: A too low GC penalty weight (1.0) results in insufficient constraint strength, a low overall GC composition; conversely, an extremely strict penalty (10.0) strictly locks in nucleotide proportions but severely restricts the exploration space of the strategy, causing a sharp drop in TE gains and undermining the original intent of generative optimization. Similarly, tests of MFE boundaries with a fixed GC weight showed that a low weight of 0.5 fails to ensure reasonable structural compactness; whereas when the weight is adjusted to just 1.2, the model begins to generate unnatural sequences containing extreme hairpin structures (with the average MFE plummeting below -31 kcal/mol). Such excessive thermodynamic stability would, in fact, severely hinder ribosomal scanning and translation initiation in real cellular environments. Following rigorous boundary testing and trade-off analysis, we ultimately finalized the optimal hyperparameter combination for driving the target-oriented generation of 5′ UTRs: a TE weight of 1.0, a GC penalty weight of 1.5, and a normalized MFE weight of 1.0. This combination provides a solid foundation that balances efficiency and high fidelity for all subsequent sequence design experiments in this paper.

### 1.4 Training Details

#### 1.4.1 Pre-training Experiment Setup

The UTRGen pre-training model adopts a GPT-2-based autoregressive architecture. The core of the network consists of 8 stacked Transformer decoders, each configured with 16 self-attention heads and a hidden dimension of 384, resulting in a total of approximately 14.3 million parameters. To fully cover the length distribution of the vast majority of natural 5′ UTRs, the maximum input sequence length of the model is set to 256 nt. During the model training phase, we use the AdamW optimizer for parameter updates, with an initial learning rate set to 3 × 10^−4^ and a CosineAnnealingLR scheduler configured for a period of 100 epochs. The pre-training process was conducted in parallel on two NVIDIA A100 GPUs with a global batch size of 512; training for 100 epochs took approximately 12 days. Since continued training led to significant overfitting, we ultimately extracted the model weights from the 100th epoch as a baseline checkpoint for subsequent evaluation and task fine-tuning.

#### 1.4.2 Experimental Setup for Downstream Function Prediction Tasks

All model fine-tuning for downstream function prediction tasks was conducted on a single NVIDIA A100 or A40 GPU. For model optimization, we uniformly employed the AdamW optimizer combined with a cosine annealing strategy (T_*max*_ = 50) for parameter updates, and the Huber loss is used as the objective function for regression tasks. For the MRL prediction task, the global batch size is set to 64, the initial learning rate is 2 × 10^−3^, and the maximum training duration is 50 epochs. For the TE, EL, and IRES prediction tasks, the batch size is also set to 64, the initial learning rate is adjusted to 2 × 10^−4^, the maximum training duration is extended to 200 epochs, and a fixed random seed of 42 is used to perform 10-fold cross-validation.

#### 1.4.3 Reinforcement Learning Fine-Tuning Experiment Setup

During the phase of RL, we fine-tuned the model using the GRPO algorithm. The entire RL training process was deployed on a single NVIDIA A100 GPU, with a total runtime of approximately 4.5 hours. The learning rate used for model optimization was set to 1 × 10^−4^. Regarding specific sampling and parameter update configurations, the global batch size was set to 32, and the group size was set to 8 (i.e., for each input prompt, 8 candidate sequences were generated for relative advantage evaluation). The model executed a total of 2,000 optimization steps, a process equivalent to an optimization scale covering approximately 512k training data points.

## 2 Supplementary Figures

**Supplementary Figure 1:**
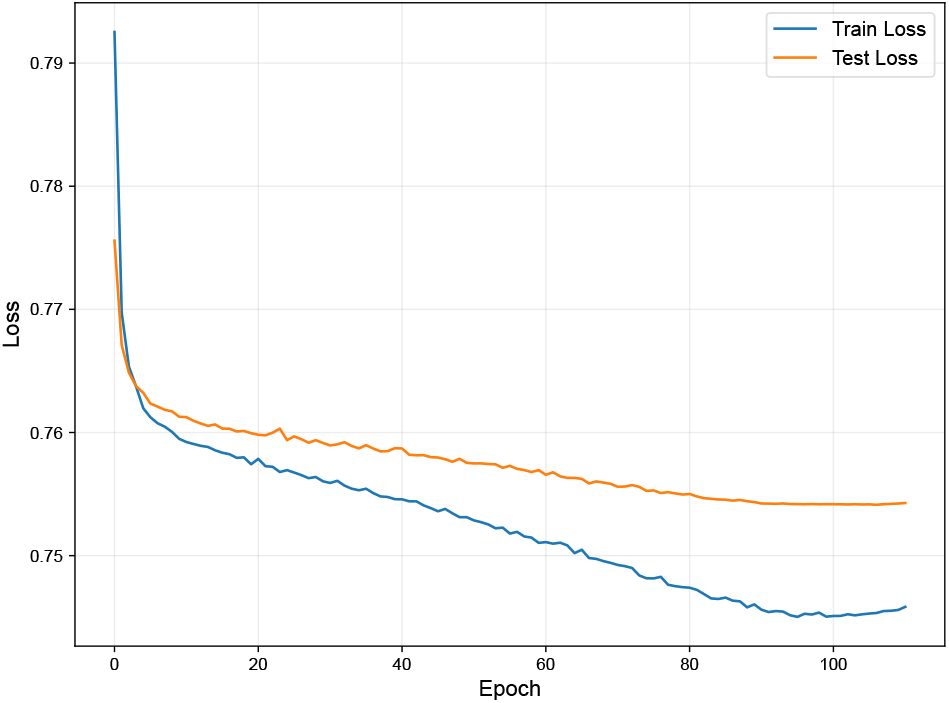
Training and testing loss curves for the UTRGen model (8 Transformer layers, 16 attention heads, 384-dimensional embeddings, maximum sequence length of 256, learning rate of 0.003) over 110 epochs. Both loss metrics declined rapidly during the first 20 epochs and then gradually leveled off. The test loss stabilized around the 90th epoch and began to diverge slightly from the training loss, indicating mild overfitting in the later stages. We used the model weights from the 100th epoch as the pre-training results.

**Supplementary Figure 2:**
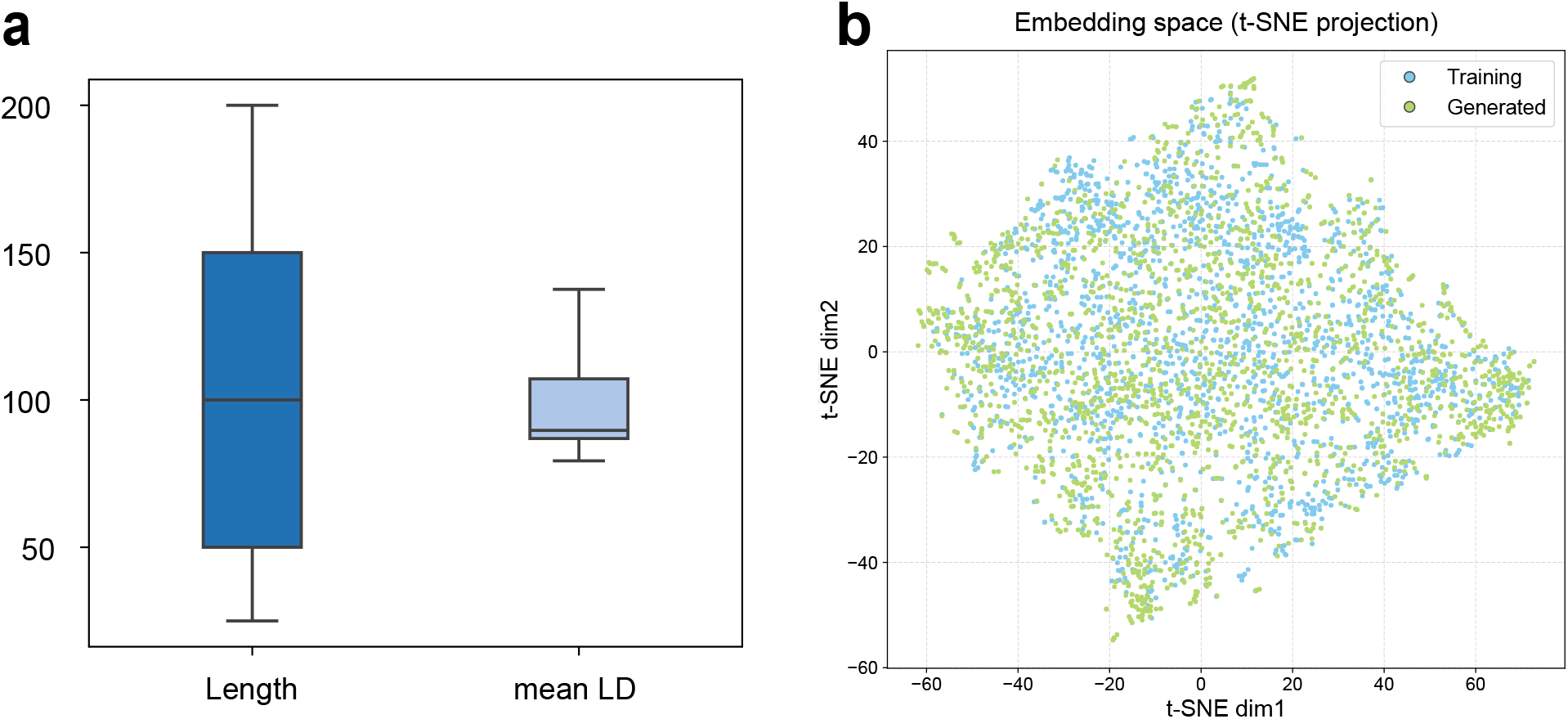
**a**, Box plots showing the length and log-average edit distance (LD) of sequences generated by UTRGen. The horizontal line within each box represents the median. **b**, Two-dimensional visualization of the sequence embedding space. A t-SNE projection of the high-dimensional hidden states extracted from the final layer of the pre-trained model. Blue points represent natural 5′ UTR sequences from the training set, while green points represent novel sequences generated by the UTRGen model. The extensive overlap between the distributions of the two groups indicates that unsupervised pre-training successfully enabled the model to capture and reproduce the underlying latent manifold of natural 5′ UTR sequences.

**Supplementary Figure 3:**
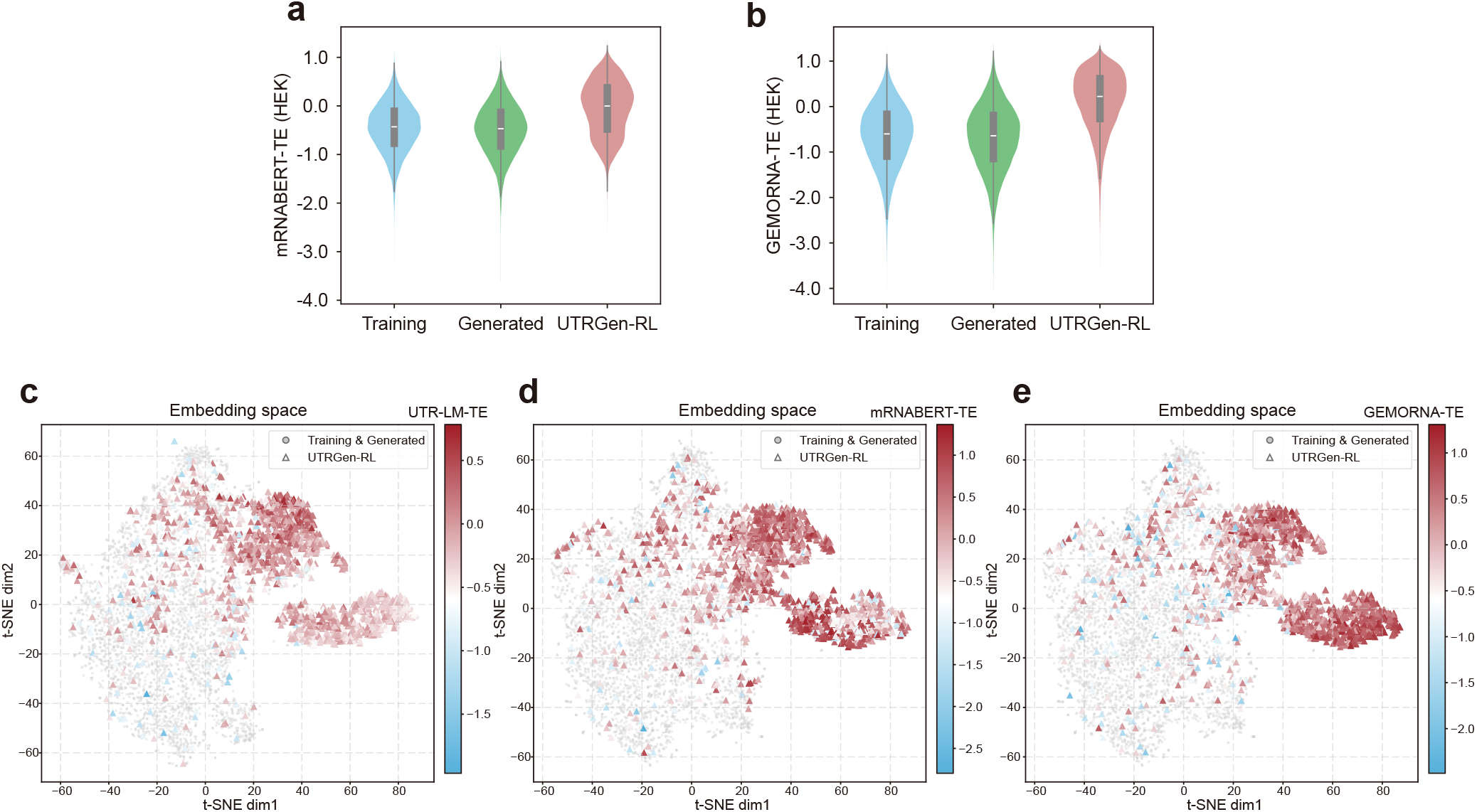
Orthogonal TE predictors further support score enrichment and spatial localization of RL-optimized sequences. **a**,**b**, Violin plots of predicted TE scores for Training, Generated, and UTRGen-RL sequences evaluated by two independent third-party predictors, GEMORNA-TE (**a**) and mRNABERT-TE (**b**). In both cases, UTRGen-RL shows an upward shift in score distribution relative to both Training and Generated sequences, consistent with the trend observed for UTR-LM-TE in Fig. 6c and for the internal reward model in Fig. 6b. **c–e**, The same t-SNE projection as in Fig. 6f,g, colored by predicted TE scores from UTR-LM-TE (**c**), GEMORNA-TE (**d**), and mRNABERT-TE (**e**). Gray circles denote Training and Generated sequences, and colored triangles denote UTRGen-RL sequences. Across all three independent predictors, the RL-enriched cluster maps to a region associated with elevated predicted TE, recapitulating the spatial pattern observed with the reward model UTRGen-TE (HEK) in Fig. 6g.

**Supplementary Figure 4:**
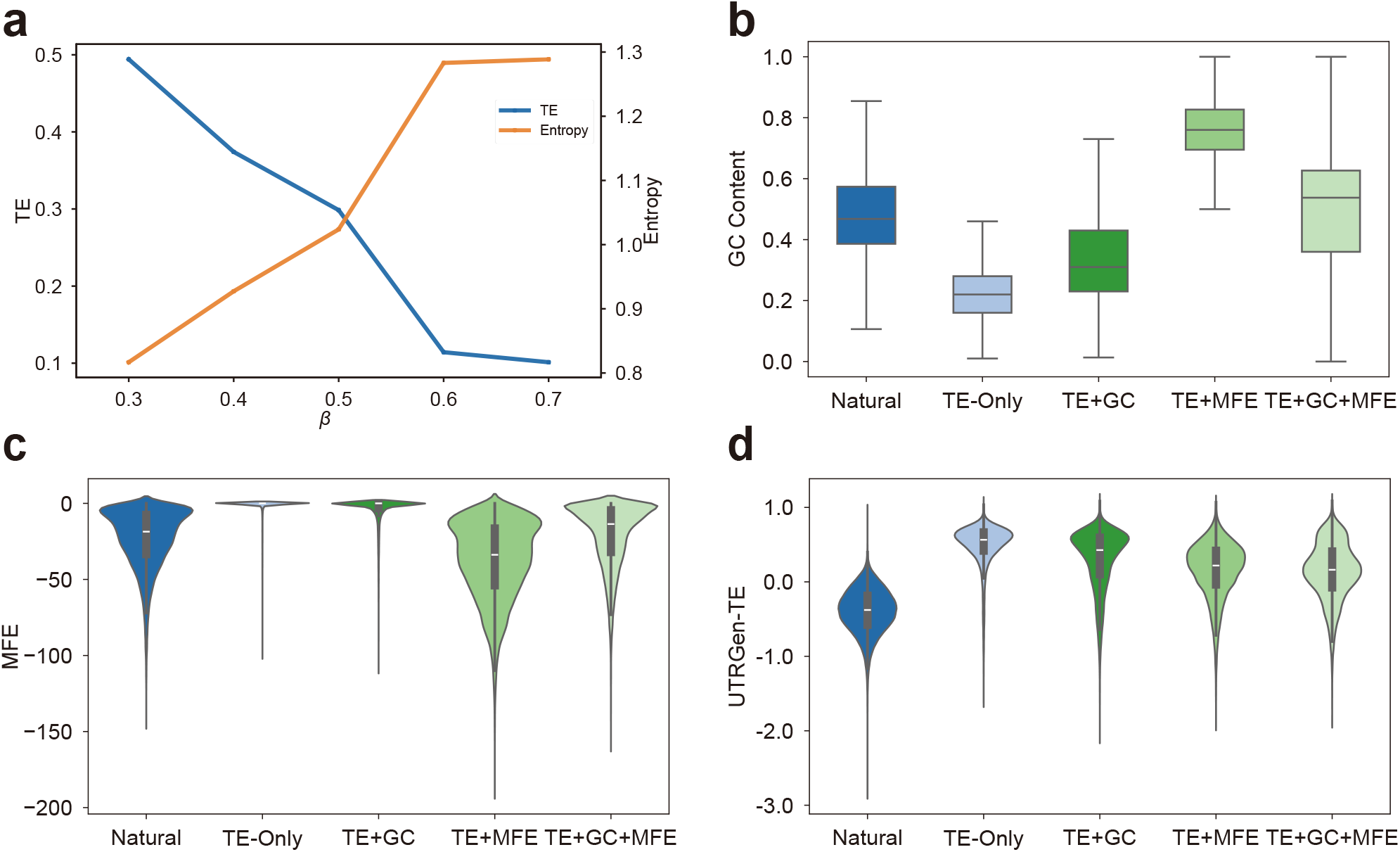
Ablation study of reinforcement learning parameters and the composite reward function. **a**, Sensitivity analysis of the KL divergence penalty coefficient (*β*) on model optimization performance. The dual-axis line chart shows the trends of predicted TE (blue line) and sequence diversity (Shannon entropy, orange line) across varying *β* values, revealing the trade-off between target optimization and sequence space exploration. **b-d**, Comparison of feature distributions of generated sequences under different reward combinations (Natural sequence baseline, TE-Only, TE+GC, TE+MFE, and the complete TE+GC+MFE reward). Shown are the box plot of GC content distribution **(b)**, the violin plot of MFE distribution **(c)**, and the violin plot of UTRGEN-TE predicted score distribution **(d)**.

**Supplementary Figure 5:**
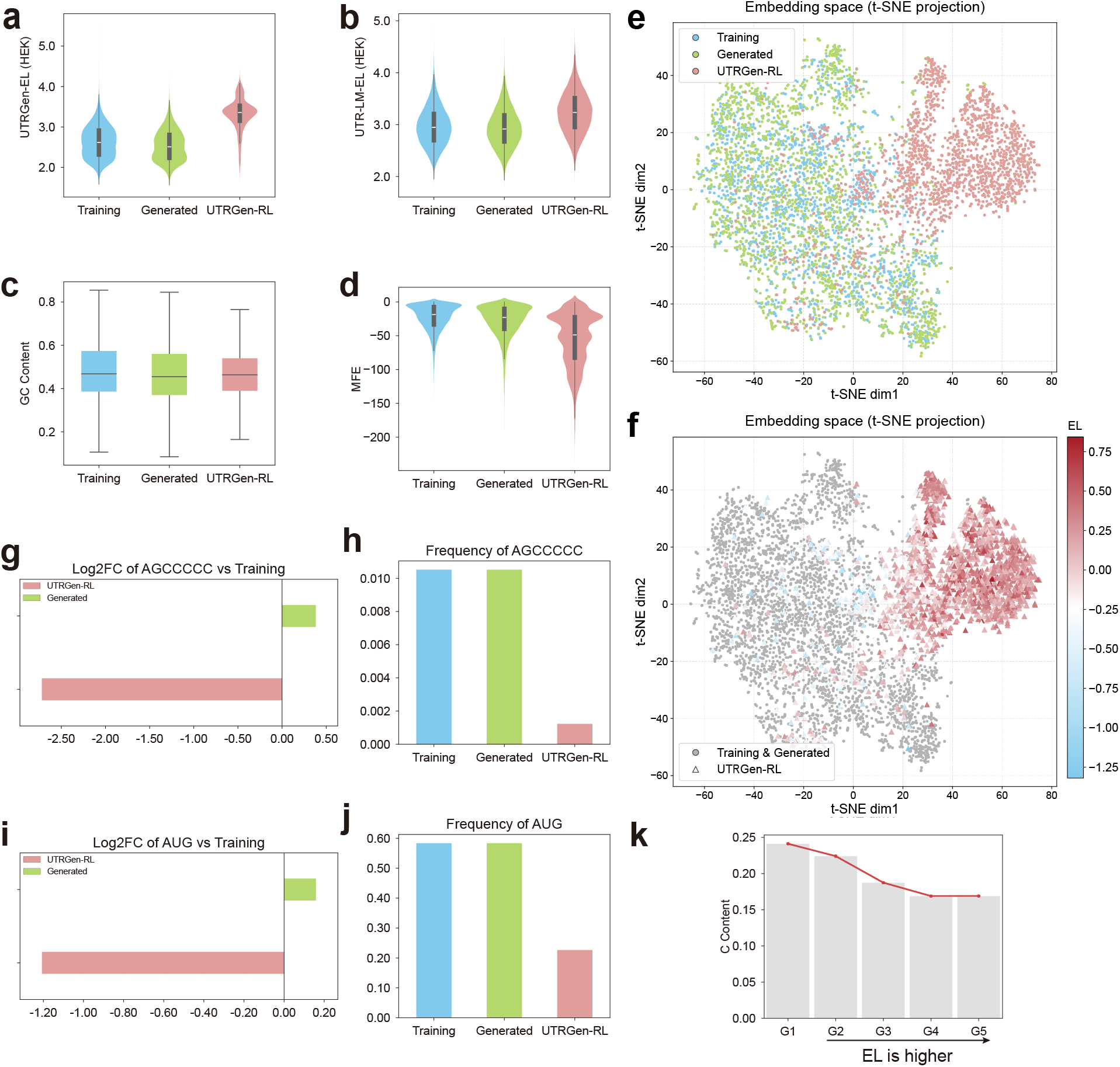
KL-regularized RL enables constraint-aware, EL-guided 5′ UTR design. **a, b**, Distributions of predicted EL scores for Training, Generated and RL-optimized sequences (UTRGen-RL), evaluated using the internal reward model UTRGen-EL (HEK) (**a**) and an independent predictor UTR-LM-EL (HEK) (**b**). **c, d**, GC content (**c**) and MFE (**d**) distributions across Training, Generated, and UTRGen-RL, showing that RL improves EL while maintaining native-like composition and secondary-structure stability. **e**, t-SNE projection of final-layer sequence embeddings for Training, Generated, and UTRGen-RL sequences, highlighting a distributional shift of RL-optimized sequences toward a distinct region in representation space. **f**, The same embedding space coloured by predicted EL, showing that the RL-enriched region coincides with high predicted EL. **g, h**, Depletion of the inhibitory motif AGCCCCC, quantified by log2FC relative to Training (**g**), and its frequency across sets (**h**). **i, j**, Depletion of uAUG, quantified by log2FC relative to Training (**i**), and its frequency across sets (**j**). **k**, Cytidine content decreases with increasing predicted EL across UTRGen-RL deciles.

**Supplementary Figure 6:**
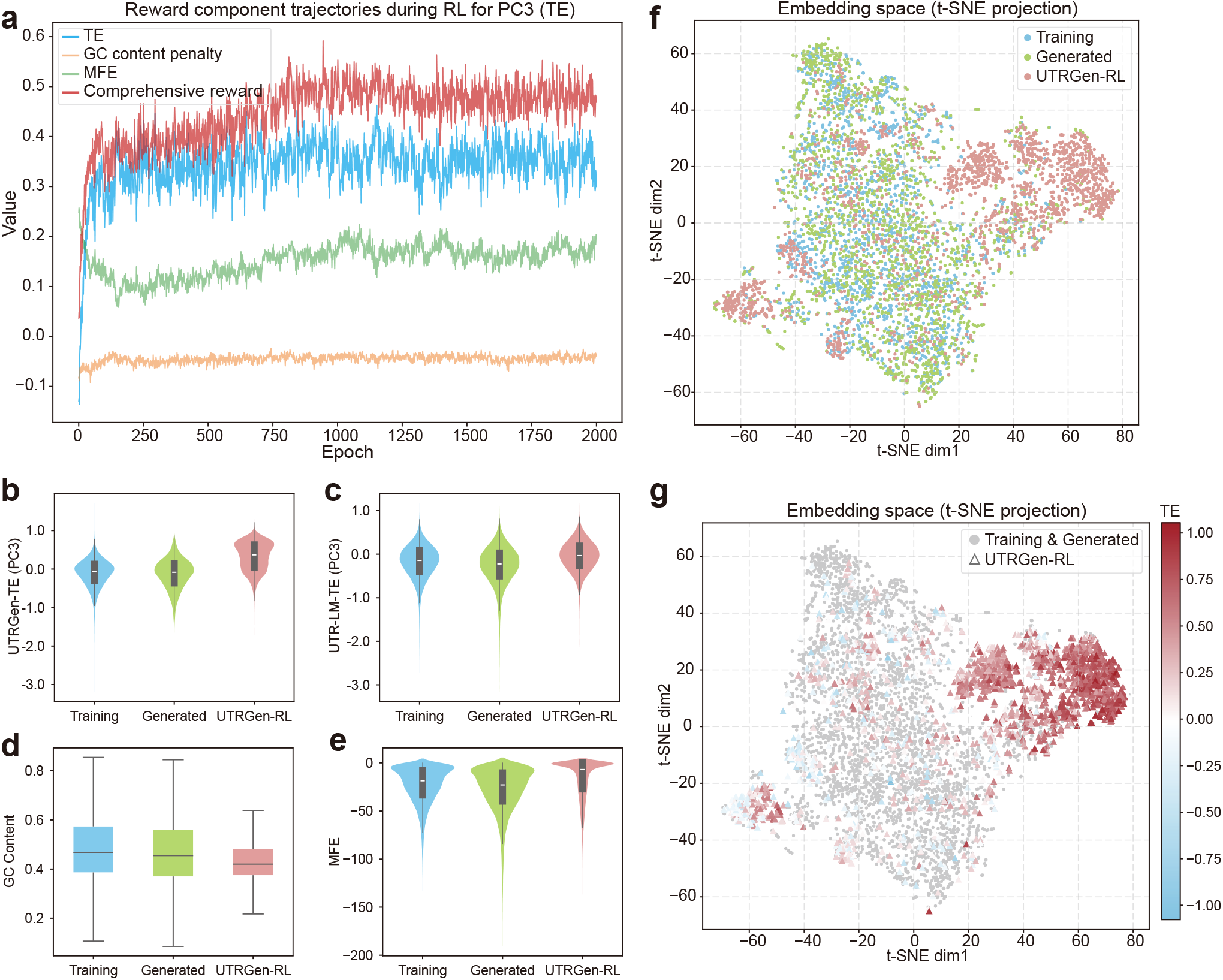
KL-regularized RL enables constraint-aware, TE-guided 5′ UTR design on PC3. **a**, Trajectories of the reward components during RL training, including the predicted TE reward, GC-content penalty, MFE-related term, and the total weighted reward, showing stable convergence under multi-objective constraints. **b, c**, Distributions of predicted TE scores for Training, Generated and RL-optimized sequences (UTRGen-RL), evaluated using the internal reward model UTRGen-TE (PC3) (**b**) and an independent predictor UTR-LM-TE (PC3) (**c**). **d, e**, GC content (**d**) and MFE (**e**) distributions across Training, Generated, and UTRGen-RL, showing that RL improves TE while maintaining native-like composition and secondary-structure stability. **f**, t-SNE projection of final-layer sequence embeddings for Training, Generated, and UTRGen-RL sequences, highlighting a distributional shift of RL-optimized sequences toward a distinct region in representation space. **g**, The same embedding space coloured by predicted TE, showing that the RL-enriched region coincides with high predicted TE.

**Supplementary Figure 7:**
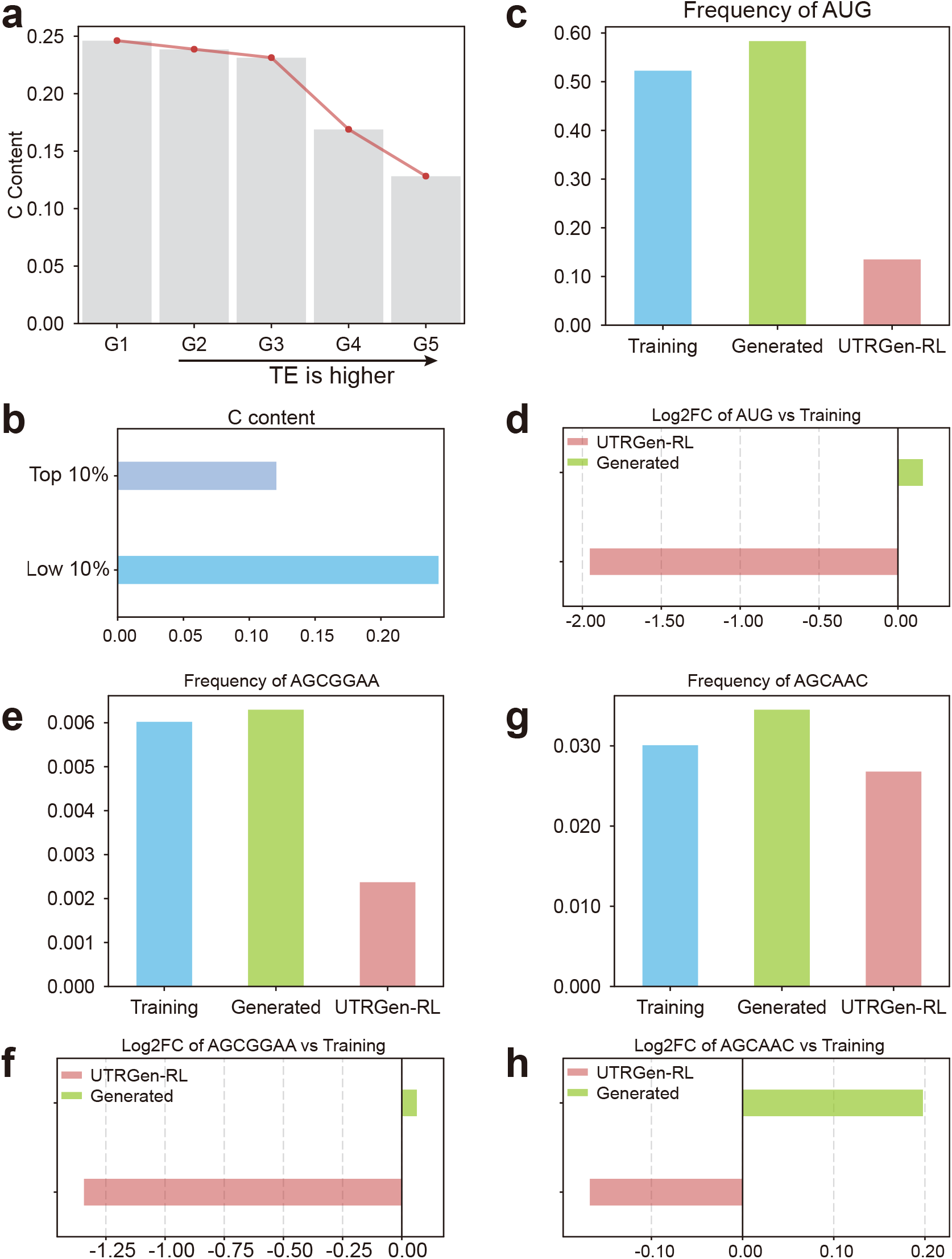
Interpreting key translation-associated features and motifs in RL-optimized 5′ UTRs on PC3. **a, b**, Cytidine content decreases with increasing predicted TE across UTRGen-RL deciles (**a**), and the top 10% UTRGen-RL sequences show lower C content than the bottom 10% (**b**). **c, d**, Frequency of uAUG across sets (**c**) and depletion quantified by log2FC relative to Training (**d**). **e-h**, Frequency of the inhibitory motif AGCGGAA (**e**) and AGCAAC (**g**) across sets and depletion quantified by log2FC relative to Training (**f**,**h**).

## 3 Supplementary Tables

**Supplementary Table 1:** Comparison of MRL prediction tasks on eight synthetic libraries with random 50 nt 5′ UTRs.

| Spearman $R$ | $U_1$ | $U_2$ | $\Psi_1$ | $\Psi_2$ | $m1\Psi_1$ | $m1\Psi_2$ | $mC-U_1$ | $mC-U_2$ |
| --- | --- | --- | --- | --- | --- | --- | --- | --- |
| UTRGen | 0.964 | 0.926 | 0.903 | 0.905 | 0.886 | 0.898 | 0.803 | 0.863 |
| mRNABERT | 0.962 | 0.924 | 0.880 | 0.901 | 0.876 | 0.862 | 0.797 | 0.787 |
| GEMORNA-UTR | 0.953 | 0.901 | 0.875 | 0.883 | 0.884 | 0.872 | 0.757 | 0.821 |
| UTR-LM | 0.962 | 0.924 | 0.898 | 0.898 | 0.875 | 0.891 | 0.772 | 0.838 |
| CaLM | 0.833 | 0.827 | 0.794 | 0.811 | 0.790 | 0.773 | 0.666 | 0.661 |
| mRNAFM | 0.819 | 0.804 | 0.758 | 0.787 | 0.771 | 0.759 | 0.644 | 0.650 |
| Optimus | 0.951 | 0.900 | 0.844 | 0.817 | 0.822 | 0.840 | 0.695 | 0.779 |
| FramePool | 0.938 | 0.898 | 0.879 | 0.879 | 0.856 | 0.863 | 0.753 | 0.779 |
| MTtrans | 0.953 | 0.912 | 0.895 | 0.892 | 0.871 | 0.882 | 0.768 | 0.807 |
| RNAErnie | 0.863 | 0.832 | 0.802 | 0.825 | 0.788 | 0.785 | 0.668 | 0.553 |
| ERNIE-RNA | 0.945 | 0.915 | 0.859 | 0.893 | 0.881 | 0.862 | 0.792 | 0.799 |
| RiNALMo | 0.931 | 0.930 | 0.867 | 0.900 | 0.870 | 0.812 | 0.776 | 0.801 |
| RNA-FM | 0.528 | 0.513 | 0.481 | 0.509 | 0.471 | 0.503 | 0.407 | 0.394 |
| RNABERT | 0.226 | 0.224 | 0.138 | 0.251 | 0.161 | 0.198 | 0.192 | 0.211 |

**Supplementary Table 2:** Comparison of MRL prediction tasks on Random and Human 5′ UTRs of varying lengths.

| Spearman $R$ | Random 5' UTRs | | | | | Human 5' UTRs | | | | |
| --- | --- | --- | --- | --- | --- | --- | --- | --- | --- | --- |
|  | 25-44 | 45-64 | 65-84 | 85-100 | All | 25-44 | 45-64 | 65-84 | 85-100 | All |
| UTRGen | 0.917 | 0.924 | 0.934 | 0.922 | 0.935 | 0.861 | 0.869 | 0.861 | 0.835 | 0.862 |
| mRNABERT | 0.834 | 0.831 | 0.794 | 0.774 | 0.815 | 0.823 | 0.816 | 0.820 | 0.802 | 0.822 |
| GEMORNA-UTR | 0.895 | 0.914 | 0.924 | 0.906 | 0.923 | 0.833 | 0.859 | 0.856 | 0.839 | 0.855 |
| UTR-LM | 0.882 | 0.904 | 0.916 | 0.886 | 0.911 | 0.823 | 0.851 | 0.848 | 0.829 | 0.846 |
| MTtrans | 0.864 | 0.889 | 0.905 | 0.891 | 0.904 | 0.822 | 0.842 | 0.837 | 0.828 | 0.840 |
| Optimus | 0.874 | 0.896 | 0.904 | 0.885 | 0.901 | 0.799 | 0.834 | 0.840 | 0.821 | 0.831 |
| RNA-FM | 0.865 | 0.884 | 0.888 | 0.868 | 0.890 | 0.798 | 0.832 | 0.838 | 0.813 | 0.830 |
| RNABERT | 0.877 | 0.880 | 0.876 | 0.847 | 0.882 | 0.795 | 0.795 | 0.780 | 0.749 | 0.783 |
| FramePool | 0.857 | 0.885 | 0.885 | 0.858 | 0.883 | 0.788 | 0.828 | 0.822 | 0.810 | 0.821 |

**Supplementary Table 3:** Comparison of TE and EL prediction (ten-fold cross-validation). Values are mean ± std.

| Spearman $R$ | TE | | | EL | | |
| --- | --- | --- | --- | --- | --- | --- |
|  | Muscle | PC3 | HEK | Muscle | PC3 | HEK |
| UTRGen | 0.772 ± 0.036 | 0.734 ± 0.016 | 0.701 ± 0.015 | 0.781 ± 0.032 | 0.726 ± 0.020 | 0.739 ± 0.011 |
| mRNABERT | 0.695 ± 0.052 | 0.683 ± 0.022 | 0.638 ± 0.021 | 0.720 ± 0.057 | 0.648 ± 0.019 | 0.653 ± 0.026 |
| GEMORNA-UTR | 0.656 ± 0.070 | 0.610 ± 0.010 | 0.561 ± 0.020 | 0.668 ± 0.050 | 0.498 ± 0.030 | 0.534 ± 0.030 |
| mRNA2Vec | 0.639 ± 0.051 | 0.621 ± 0.026 | 0.591 ± 0.027 | 0.655 ± 0.051 | 0.582 ± 0.022 | 0.612 ± 0.034 |
| UTR-LM | 0.670 ± 0.040 | 0.650 ± 0.020 | 0.600 ± 0.020 | 0.660 ± 0.060 | 0.630 ± 0.020 | 0.650 ± 0.020 |
| Optimus | 0.410 ± 0.110 | 0.380 ± 0.090 | 0.360 ± 0.080 | 0.150 ± 0.110 | 0.190 ± 0.060 | 0.180 ± 0.050 |
| Cao-RF | 0.630 ± 0.040 | 0.630 ± 0.020 | 0.550 ± 0.020 | 0.640 ± 0.040 | 0.590 ± 0.030 | 0.570 ± 0.020 |
| FramePool | 0.310 ± 0.133 | 0.340 ± 0.064 | 0.350 ± 0.043 | 0.280 ± 0.085 | 0.330 ± 0.037 | 0.370 ± 0.050 |
| MTtrans | 0.500 ± 0.068 | 0.540 ± 0.023 | 0.500 ± 0.016 | 0.480 ± 0.065 | 0.480 ± 0.036 | 0.470 ± 0.021 |
| RNA-FM | 0.300 ± 0.058 | 0.320 ± 0.030 | 0.230 ± 0.024 | 0.300 ± 0.061 | 0.200 ± 0.014 | 0.220 ± 0.025 |
| RNABERT | 0.560 ± 0.060 | 0.480 ± 0.040 | 0.410 ± 0.040 | 0.600 ± 0.040 | 0.450 ± 0.060 | 0.470 ± 0.050 |

**Supplementary Table 4:** Statistical significance of benchmark comparisons for TE and EL prediction. *P* values were computed using paired *t*-tests on fold-wise Spearman *R* values from ten-fold cross-validation, comparing UTRGen with each baseline method.

| $P$ value | TE | | | EL | | |
| --- | --- | --- | --- | --- | --- | --- |
|  | Muscle | PC3 | HEK | Muscle | PC3 | HEK |
| mRNABERT | 0.0014 | $1.63 \times 10^{-5}$ | $9.51 \times 10^{-7}$ | 0.0104 | $5.80 \times 10^{-8}$ | $4.63 \times 10^{-7}$ |
| GEMORNA-UTR | $4.00 \times 10^{-4}$ | $1.60 \times 10^{-12}$ | $2.98 \times 10^{-12}$ | $2.15 \times 10^{-5}$ | $1.40 \times 10^{-12}$ | $2.73 \times 10^{-10}$ |
| mRNA2Vec | $4.49 \times 10^{-6}$ | $6.24 \times 10^{-9}$ | $1.99 \times 10^{-8}$ | $7.82 \times 10^{-6}$ | $1.05 \times 10^{-11}$ | $2.55 \times 10^{-7}$ |
| UTR-LM | $1.20 \times 10^{-5}$ | $8.18 \times 10^{-9}$ | $4.85 \times 10^{-10}$ | $6.70 \times 10^{-5}$ | $2.97 \times 10^{-9}$ | $6.66 \times 10^{-9}$ |
| Optimus | $8.84 \times 10^{-7}$ | $3.67 \times 10^{-7}$ | $1.68 \times 10^{-7}$ | $4.30 \times 10^{-9}$ | $2.36 \times 10^{-11}$ | $1.27 \times 10^{-11}$ |
| Cao-RF | $1.45 \times 10^{-7}$ | $3.11 \times 10^{-10}$ | $8.93 \times 10^{-13}$ | $1.05 \times 10^{-7}$ | $2.83 \times 10^{-9}$ | $1.28 \times 10^{-12}$ |
| FramePool | $7.16 \times 10^{-7}$ | $3.19 \times 10^{-9}$ | $4.96 \times 10^{-11}$ | $1.25 \times 10^{-9}$ | $5.95 \times 10^{-14}$ | $7.29 \times 10^{-10}$ |
| MTtrans | $2.93 \times 10^{-8}$ | $2.19 \times 10^{-13}$ | $1.65 \times 10^{-16}$ | $6.28 \times 10^{-9}$ | $2.14 \times 10^{-11}$ | $7.33 \times 10^{-15}$ |
| RNA-FM | $8.22 \times 10^{-13}$ | $2.16 \times 10^{-15}$ | $1.58 \times 10^{-18}$ | $4.76 \times 10^{-12}$ | $2.92 \times 10^{-21}$ | $1.27 \times 10^{-16}$ |
| RNABERT | $1.02 \times 10^{-7}$ | $4.02 \times 10^{-10}$ | $1.21 \times 10^{-10}$ | $2.66 \times 10^{-9}$ | $2.80 \times 10^{-8}$ | $1.53 \times 10^{-8}$ |

**Supplementary Table 5:** Performance on the IRES dataset. Values are mean ± std (ten-fold cross-validation).

| Method | ACC | AUC | AUPR |
| --- | --- | --- | --- |
| UTRGen | $0.639 \pm 0.007$ | $0.685 \pm 0.009$ | $0.715 \pm 0.010$ |
| mRNABERT | $0.626 \pm 0.007$ | $0.677 \pm 0.006$ | $0.712 \pm 0.006$ |
| UTR-LM | $0.618 \pm 0.005$ | $0.660 \pm 0.009$ | $0.686 \pm 0.011$ |
| GEMORNA-UTR | $0.602 \pm 0.007$ | $0.640 \pm 0.011$ | $0.668 \pm 0.012$ |
| IRESpy | $0.620 \pm 0.007$ | $0.667 \pm 0.008$ | $0.699 \pm 0.009$ |
| FramePool | $0.615 \pm 0.007$ | $0.662 \pm 0.006$ | $0.693 \pm 0.008$ |
| IRESfinder | $0.569 \pm 0.006$ | $0.599 \pm 0.007$ | $0.591 \pm 0.010$ |

**Supplementary Table 6:** Statistical significance of benchmark comparisons for IRES prediction. *P* values were computed using paired *t*-tests on fold-wise results from ten-fold cross-validation, comparing UTRGen with each baseline method. Values marked as ns are not statistically significant (*P* ≥ 0.05).

| $P$ value | ACC | AUC | AUPR |
| --- | --- | --- | --- |
| mRNABERT | $6.00 \times 10^{-4}$ | 0.0329 | 0.4289<sup>ns</sup> |
| UTR-LM | $7.82 \times 10^{-7}$ | $7.33 \times 10^{-6}$ | $8.32 \times 10^{-6}$ |
| GEMORNA-UTR | $6.44 \times 10^{-10}$ | $1.27 \times 10^{-8}$ | $2.55 \times 10^{-8}$ |
| IRESpy | $9.78 \times 10^{-6}$ | $1.74 \times 10^{-4}$ | 0.0015 |
| FramePool | $4.47 \times 10^{-7}$ | $5.42 \times 10^{-6}$ | $4.33 \times 10^{-5}$ |
| IRESfinder | $6.86 \times 10^{-15}$ | $1.72 \times 10^{-14}$ | $3.22 \times 10^{-16}$ |

**Supplementary Table 7:** Ten 5′ UTR sequence libraries used for MRL prediction. Synthetic libraries (U_1_, U_2_, mC-U_1_, mC-U_2_, Ψ_1_, Ψ_2_, m1Ψ_1_, and m1Ψ_2_) contain fixed-length (50nt) sequences, while the “Random” and “Human” datasets include variable-length sequences.

| Library | CDS | Uridine | Counts (#) | Length |
| --- | --- | --- | --- | --- |
| $U_1$ | eGFP | Unmodified | 280,000 | 50 |
| $U_2$ | eGFP | Unmodified | 280,000 | 50 |
| mC-$U_1$ | mCherry | Unmodified | 280,000 | 50 |
| mC-$U_2$ | mCherry | Unmodified | 280,000 | 50 |
| $\Psi_1$ | eGFP | Pseudouridine | 279,968 | 50 |
| $\Psi_2$ | eGFP | Pseudouridine | 280,000 | 50 |
| m1$\Psi_1$ | eGFP | 1-methylpseudouridine | 279,968 | 50 |
| m1$\Psi_2$ | eGFP | 1-methylpseudouridine | 280,000 | 50 |
| Random | - | - | 83,919 | 25-100 |
| Human | - | - | 15,555 | 25-100 |

**Supplementary Table 8:** Summary of three datasets of human endogenous 5′ UTRs with mRNA TE and EL.

| Source | Counts (#) | Mean of Length | Median of Length |
| --- | --- | --- | --- |
| Muscle | 1257 | 183 | 126 |
| PC3 | 12,579 | 192 | 132 |
| HEK | 14,410 | 185 | 127 |

**Supplementary Table 9:** Numbers of IRES and non-IRES sequences in each length range.

| Length range | Non-IRES | IRES | Total |
| --- | --- | --- | --- |
| < 174 | 53 | 991 | 1044 |
| 174 | 25534 | 21107 | 46641 |
| 175-200 | 3 | 592 | 595 |
| 200-300 | 27 | 1223 | 1250 |
| 300-500 | 86 | 1002 | 1088 |
| 500-700 | 35 | 226 | 261 |
| 700-1000 | 51 | 45 | 96 |
| 1000-2000 | 54 | 24 | 78 |
| Total | 25843 | 25210 | 51053 |

**Supplementary Table 10:** Reward Weight Analysis Results.

| $w_{TE}$ | $w_{GC\text{penalty}}$ | $w_{MFE}$ | TE | GC Content | MFE |
| --- | --- | --- | --- | --- | --- |
| 1 | 1 | 1 | 0.1752 | 0.4443 | -14.9507 |
| **1** | **1.5** | **1** | **0.1413** | **0.4982** | **-20.7686** |
| 1 | 2 | 1 | 1.1331 | 0.514 | -21.9567 |
| 1 | 10 | 1 | 0.0743 | 0.5458 | -24.7347 |
| 1 | 1.5 | 0.5 | 0.0976 | 0.4475 | -14.6267 |
| 1 | 1.5 | 1.2 | 0.0542 | 0.5883 | -32.4555 |

## References

[1] Eduarde Rohner, Ran Yang, Kylie S Foo, Alexander Goedel, and Kenneth R Chien. Unlocking the promise of mrna therapeutics. Nature biotechnology, 40(11):1586–1600, 2022.

[2] Patricia R Araujo, Kihoon Yoon, Daijin Ko, Andrew D Smith, Mei Qiao, Uthra Suresh, Suzanne C Burns, and Luiz OF Penalva. Before it gets started: regulating translation at the 5′ utr. International Journal of Genomics, 2012(1):475731, 2012.

[3] Alan G Hinnebusch, Ivaylo P Ivanov, and Nahum Sonenberg. Translational control by 5′-untranslated regions of eukaryotic mrnas. Science, 352(6292):1413–1416, 2016.

[4] Kathrin Leppek, Rhiju Das, and Maria Barna. Functional 5′ utr mrna structures in eukaryotic translation regulation and how to find them. Nature reviews Molecular cell biology, 19(3):158–174, 2018.

[5] Paul J Sample, Ban Wang, David W Reid, Vlad Presnyak, Iain J McFadyen, David R Morris, and Georg Seelig. Human 5′ utr design and variant effect prediction from a massively parallel translation assay. Nature biotechnology, 37(7):803–809, 2019.

[6] Jicong Cao, Eva Maria Novoa, Zhizhuo Zhang, William CW Chen, Dianbo Liu, Gigi CG Choi, Alan SL Wong, Claudia Wehrspaun, Manolis Kellis, and Timothy K Lu. High-throughput 5′ utr engineering for enhanced protein production in non-viral gene therapies. Nature communications, 1(12):4138, 2021.

[7] Weizhong Zheng, John HC Fong, Yuk Kei Wan, Athena HY Chu, Yuanhua Huang, Alan SL Wong, and Joshua WK Ho. Discovery of regulatory motifs in 5′ untranslated regions using interpretable multi-task learning models. Cell Systems, 14(12):1103–1112, 2023.

[8] Alexander Karollus, Žiga Avsec, and Julien Gagneur. Predicting mean ribosome load for 5′ utr of any length using deep learning. PLoS computational biology, 5(17):e1008982, 2021.

[9] Yanyi Chu, D. Yin Dan Yu, Guangxue Xu, Junze Zhang, Xiaotong Wang, Yue Shen, Yupeng Li, Ning Zhao, Yi Zhu, et al. Programmable rna translation through deep learning-driven ires discovery and de novo generation. Nature Machine Intelligence, 8(4):559–574, 2026.

[10] Yanyi Chu, Dan Yu, Yupeng Li, Kaixuan Huang, Yue Shen, L. Cong, Jason Zhang, and Mengdi Wang. A 5′ utr language model for decoding untranslated regions of mrna and function predictions. Nature Machine Intelligence, 6(4):449–460, 2024.

[11] Ying Xiong, Aowen Wang, Yu Kang, Chao Shen, Chang-Yu Hsieh, and Tingjun Hou. mrnabert: advancing mrna sequence design with a universal language model and comprehensive dataset. Nature Communications, 1(16):10371, 2025.

[12] He Zhang, Hailong Liu, Yushan Xu, Haoran Huang, Yiming Liu, Jia Wang, Yan Qin, Haiyan Wang, Lili Ma, Zhiyuan Xun, et al. Deep generative models design mrna sequences with enhanced translational capacity and stability. Science, 6773(390):eadr8470, 2025.

[13] Zhihong Shao, Peiyi Wang, Qihao Zhu, Runxin Xu, Junxiao Song, Xiao Bi, Haowei Zhang, Mingchuan Zhang, YK Li, Yang Wu, et al. Deepseekmath: Pushing the limits of mathematical reasoning in open language models. arXiv preprint arXiv:2402.03300, 2024.

[14] Daya Guo, Dejian Yang, Haowei Zhang, Junxiao Song, Peiyi Wang, Qihao Zhu, Runxin Xu, Ruoyu Zhang, Shirong Ma, Xiao Bi, et al. Deepseek-r1 incentivizes reasoning in llms through reinforcement learning. Nature, 645(8081):633–638, 2025.

[15] Cole JT Lewis, Li H Xie, Shivani Milind Bhandarkar, Danni Jin, Kyrillos Abdallah, Austin S Draycott, Yixuan Chen, Carson C Thoreen, and Wendy V Gilbert. Quantitative profiling of human translation initiation reveals elements that potently regulate endogenous and therapeutically modified mrnas. Molecular Cell, 85(2):445–459, 2025.

[16] Michele Iacono, Flavio Mignone, and Graziano Pesole. uaug and uorfs in human and rodent 5′ untranslated mrnas. Gene, 349:97–105, 2005.

[17] Basak Eraslan, Dongxue Wang, Mirjana Gusic, Holger Prokisch, Björn M Hallström, Mathias Uhlén, Anna Asplund, Frederik Pontén, Thomas Wieland, Thomas Hopf, et al. Quantification and discovery of sequence determinants of protein-per-mrna amount in 29 human tissues. Molecular systems biology, 2(15):MSB188513, 2019.

[18] Claudio Lo Giudice, Federico Zambelli, Matteo Chiara, Giulio Pavesi, Marco Antonio Tangaro, Ernesto Picardi, and Graziano Pesole. Utrdb 2.0: a comprehensive, expert curated catalog of eukaryotic mrnas untranslated regions. Nucleic acids research, 51(D1):D337–D344, 2023.

[19] Tamara Goldfarb, Vamsi K Kodali, Shashikant Pujar, Vyacheslav Brover, Barbara Robbertse, Catherine M Farrell, Dong-Ha Oh, Alexander Astashyn, Olga Ermolaeva, Diana Haddad, et al. Ncbi refseq: reference sequence standards through 25 years of curation and annotation. Nucleic acids research, 53(D1):D243–D257, 2025.

[20] Manato Akiyama and Yasubumi Sakakibara. Informative rna base embedding for rna structural alignment and clustering by deep representation learning. NAR genomics and bioinformatics, 1(4):qac012, 2022.

[21] Jiayang Chen, Zhihang Hu, Siqi Sun, Qingxiong Tan, Yixuan Wang, Qinze Yu, Licheng Zong, Liang Hong, Jin Xiao, Tao Shen, et al. Interpretable rna foundation model from unannotated data for highly accurate rna structure and function predictions. arXiv preprint arXiv:2204.00300, 2022.

[22] Ning Wang, Jiang Bian, Yuchen Li, Xuhong Li, Shahid Mumtaz, Linghe Kong, and Haoyi Xiong. Multi-purpose rna language modelling with motif-aware pretraining and type-guided fine-tuning. Nature Machine Intelligence, 6(5):548–557, 2024.

[23] Weijie Yin, Zhaoyu Zhang, Shuo Zhang, Liang He, Ruiyang Zhang, Rui Jiang, Gan Liu, Jingyi Wang, Xuegong Zhang, Tao Qin, et al. Ernie-rna: an rna language model with structure-enhanced representations. Nature Communications, 1(16):10076, 2025.

[24] Rafael Josip Penić, Tin Vlašić, Roland G Huber, Yue Wan, and Mile Šikić. Rinalmo: General-purpose rna language models can generalize well on structure prediction tasks. Nature Communications, 1(16):5671, 2025.

[25] Carlos Outeiral and Charlotte M Deane. Codon language embeddings provide strong signals for use in protein engineering. Nature Machine Intelligence, 6(2):170–179, 2024.

[26] Honggen Zhang, Xiangrui Gao, June Zhang, and Lipeng Lai. mrna2vec: mrna embedding with language model in the 5′ utr-cds for mrna design. In Proceedings of the AAAI Conference on Artificial Intelligence, volume 39, pages 1057–1065, 2025.

[27] Junhui Wang and Michael Gribskov. Irespy: an xgboost model for prediction of internal ribosome entry sites. BMC bioinformatics, 1(20):409, 2019.

[28] Jian Zhao, Jing Wu, Tianyi Xu, Qichang Yang, Junhao He, and Xiaofeng Song. Iresfinder: Identifying rna internal ribosome entry site in eukaryotic cell using framed k-mer features. Journal of Genetics and Genomics, 45(7):403–406, 2018.

[29] W James Kent. Blat—the blast-like alignment tool. Genome research, 12(4):656–664, 2002.

[30] Le Van Vinh, Tran Van Lang, Le Thanh Binh, and Tran Van Hoai. A two-phase binning algorithm using l-mer frequency on groups of non-overlapping reads. Algorithms for Molecular Biology, 1(10):2, 2015.

[31] Pandurang Kolekar, Abhijeet Pataskar, Urmila Kulkarni-Kale, Jayanta Pal, and Abhijeet Kulkarni. Irespred: web server for prediction of cellular and viral internal ribosome entry site (ires). Scientific reports, 1(6):27436, 2016.

[32] Jian Zhao, Yan Li, Cong Wang, Haotian Zhang, Hao Zhang, Bin Jiang Xuejiang Guo, and Xiaofeng Song. Iresbase: a comprehensive database of experimentally validated internal ribosome entry sites. Genomics, proteomics & bioinformatics, 18(2):129–139, 2020.

[33] Nancy Ontiveros-Palacios, Emma Cooke, Eric P Nawrocki, Sandra Triebel, Manja Marz, Elena Rivas, Sam Griffiths-Jones, Anton I Petrov, Alex Bateman, and Blake Sweeney. Rfam 15: Rna families database in 2025. Nucleic acids research, 53(D1):D258–D267, 2025.

[34] Martin Mokrejš, Tomáš Mašek, Václav Vopálenský, Petr Hlubuček, Philippe Delbos, and Martin Pospíšek. Iresite—a tool for the examination of viral and cellular internal ribosome entry sites. Nucleic acids research, 38(suppl_1):D131–D136, 2010.

[35] Shira Weingarten-Gabbay, Shani Elias-Kirma, Ronit Nir, Alexey A Gritsenko, Noam Stern-Ginossar, Zohar Yakhini, Adina Weinberger, and Eran Segal. Systematic discovery of cap-independent translation sequences in human and viral genomes. Science, 6270(351):aad4939, 2016.

[36] Chun-Kan Chen, Ran Cheng, Janos Demeter, Jin Chen, Shira Weingarten-Gabbay, Lihua Jiang, Michael P Snyder, Jonathan S Weissman, Eran Segal, Peter K Jackson, et al. Structured elements drive extensive circular rna translation. Molecular cell, 81(20):4300–4318, 2021.

[37] Zihe Liu, Jiashun Liu, Yancheng He, Weixun Wang, Jiaheng Liu, Ling Pan, Xinyu Hu, Shaopan Xiong, Ju Huang, Jian Hu, Shengyi Huang, Siran Yang, Jiamang Wang, Wenbo Su, and Bo Zheng. Tricks or traps? a deep dive into RL for LLM reasoning. In The Fourteenth International Conference on Learning Representations, 2026.

[38] Ronny Lorenz, Stephan H Bernhart, Christian Höner zu Siederdissen, Hakim Tafer, Christoph Flamm, Peter F Stadler, and Ivo L Hofacker. Viennarna package 2.0. Algorithms for molecular biology, 6:1–14, 2011.

[39] Martin Steinegger and Johannes Söding. Mmseqs2 enables sensitive protein sequence searching for the analysis of massive data sets. Nature biotechnology, 35(11):1026–1028, 2017.

